# Pediatric Long COVID Is Characterized by Myeloid CCR6 Suppression and Immune Dysregulation

**DOI:** 10.1101/2025.08.22.671713

**Authors:** Jon Izquierdo-Pujol, Núria Pedreño-López, Tetyana Pidkova, Maria Nevot, Victor Urrea, Fernando Laguía, Francisco Muñoz-López, Judith Dalmau, Alba Gonzalez-Aumatell, Clara Carreras-Abad, Maria Mendez, Carlos Rodrigo, Marta Massanella, Julià Blanco, Jorge Carrillo, Benjamin Trinité, Javier Martinez-Picado, Sara Morón-López

**Author notes:** Corresponding authors: Javier Martinez-Picado and Sara Morón-López.

## Abstract

The biological mechanisms underlying long COVID in the pediatric population are poorly understood. Our study aimed to characterize the immune pathophysiology of long COVID in children and young people (CYP). We analyzed major immune cell compartments in PBMCs, as well as specific SARS-CoV-2 antibody response in CYP with (n=99) and without (n=18) long COVID at three months following acute infection. Our findings indicate that pediatric long COVID is associated with a dysregulated immune response characterized by altered innate immunity and overactivated T-, B- and NK-cell responses. Furthermore, CYP with long COVID had an impaired humoral response to SARS-CoV-2 marked by a dysregulated B-cell compartment and lower levels of anti-RBD IgG and IgA. This correlated with reduced neutralizing capacity against SARS-CoV-2. Random forest analysis identified CCR6 expression on myeloid cells as the most relevant biomarker that distinguishes long COVID from control individuals with 79% accuracy.

**GRAPHICAL ABSTRACT:** 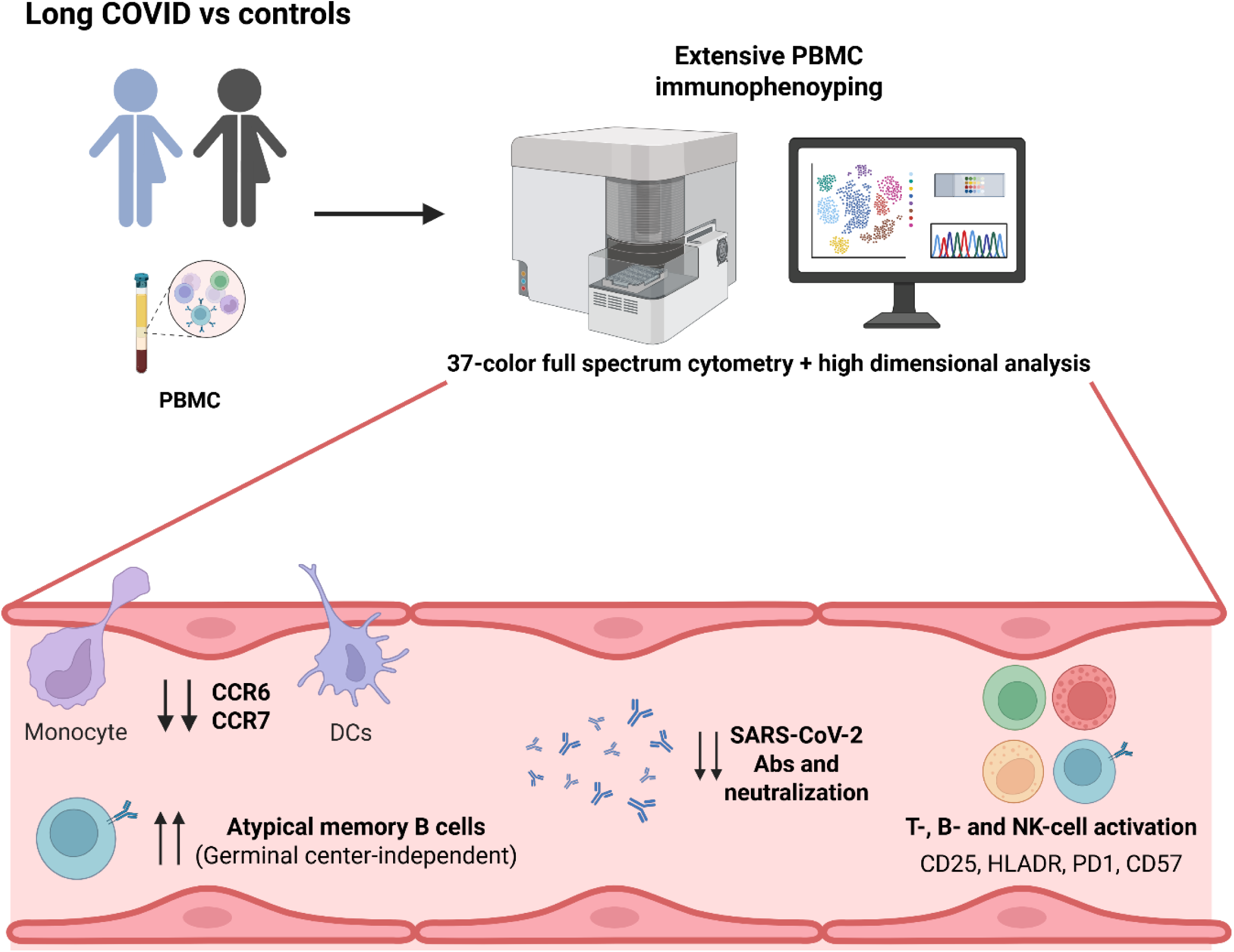

## INTRODUCTION

Long COVID (LC) affects 10-20% of people following acute SARS-CoV-2 infection^1,2^. This post-viral syndrome is not limited to adults but has also been reported in children and young people (CYP)^3–6^.

Recent research on LC has focused primarily on unraveling its underlying pathophysiologic mechanisms. Studies have demonstrated associations with immune dysregulation^7,8^, autoimmunity^9,10^, SARS-CoV-2 antigen persistence^11,12^, and viral reactivation^9,13^. However, knowledge of this condition in pediatric individuals remains limited, as most LC studies focus on the adult population, with minimal attention given to understanding the potential driving mechanisms in pediatric patients^14^.

To address this gap, we characterized the immune profile in CYP with and without LC through deep immunophenotyping of peripheral blood mononuclear cells (PBMCs) using a multiparametric 37-color spectral cytometry panel and analysis of antibody levels and ability to neutralize SARS-CoV-2.

## MATERIALS AND METHODS

### Study participants, sample processing and data collection

PediaCOVID cohort participants were recruited between January 2021 and February 2022 at University Hospital Germans Trias i Pujol. The LC cohort included individuals aged <18 years diagnosed with SARS-CoV-2 infection (PCR, antigen test and/or serology) and with at least 12 weeks of persistent COVID-19 symptoms after acute COVID-19^6^. The control cohort included individuals aged <18 years without LC, infection or immune system involvement at sample collection who had previously been diagnosed or not with SARS-CoV-2 infection (PCR, antigen test and/or serology). Parents/legal guardians and CYP older than 12 years agreed to collection of their samples and signed the informed consent document. Blood samples were prospectively collected from the pediaCOVID cohort (Hospital Ethics Committee ID #PI-21-029), and plasma and PBMCs were isolated using Ficoll-Paque and cryopreserved for further studies. We analyzed 99 samples from CYP with LC (LC cohort) and 18 from CYP without LC (control cohort). All the participants had mild or asymptomatic acute infection, were not hospitalized and did not receive any treatment or intervention during acute illness. Individuals who were infected and vaccinated received the full immunization schedule after acute infection (Two exposures to SARS-CoV-2 antigens group).

The data recorded included demographic data, medical and family history, SARS-CoV- 2 infection data, symptoms during the acute phase of COVID-19 (in symptomatic cases) and persistent symptoms (e.g. headache, fatigue, brain fog). CYP whose SARS-CoV-2 infection was not microbiologically confirmed by PCR, antigen test or serology underwent an immune functional study (SARS-CoV-2 specific T-cell immunity).

### Multiparametric spectral cytometry immunophenotyping (37-color panel)

PBMCs were stained with the 37-color antibody panel summarized in Supplementary Table 1, fixed with 1% paraformaldehyde (Sigma Aldrich) and analyzed in a Cytek Aurora 5-laser Spectral Cytometer (Cytek Biosciences). Cytometry data were analyzed using FlowJo™ v10.10 (BD Life Sciences) and OMIQ (Dotmatics: www.omiq.ai, www.dotmatics.com). A manual gating strategy was followed to export the different immune cell populations (myeloid cells, NK cells, T cells and B cells) (Supp.Fig.1). The files containing the immune cell populations were then uploaded and analyzed in the OMIQ platform. The FlowSOM algorithm was used to identify clusters of the different immune cell populations. Myeloid cells were clustered using CD1c, CD11b, CD16, CD11c, HLADR, CD95, CD14 and CD123. NK cells were clustered using CD56, CD16, CD57, NKG2A, NKG2C, NKp46, CCR5 and CXCR3. T cells were clustered using TCRVd2, CD4, CD8, CD45RA, CCR7, CD95, CD27, CD28, CCR6, CXCR3, CD25 and CD127. B cells were clustered using IgD, IgM, IgA, CD27, CD21, CD38, CD24, CD11c, CCR7, CCR6, CXCR5 and CXCR3.

**Figure 1:**
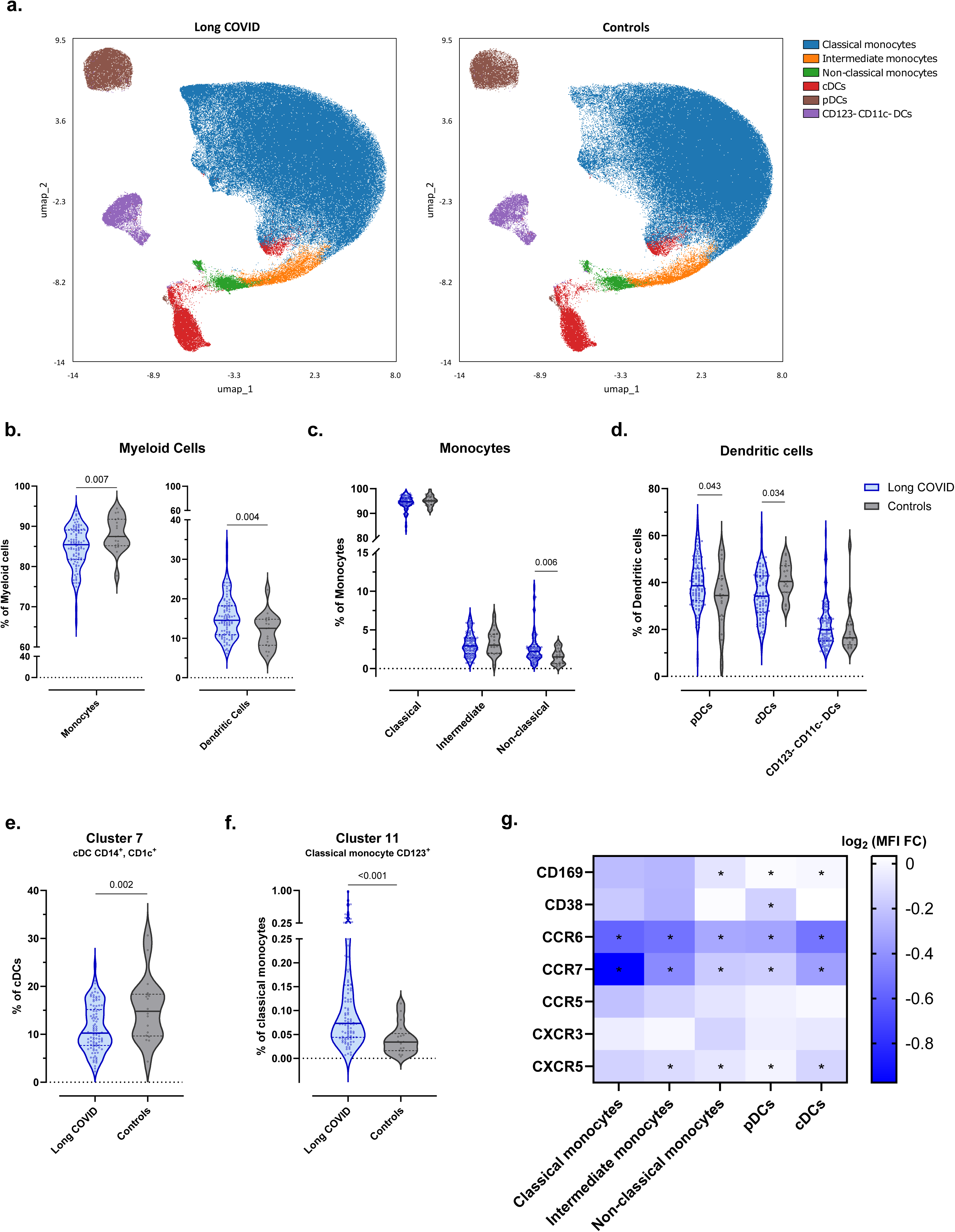
Myeloid cell compartment in CYP with and without LC. (A) Uniform manifold approximation and projection for dimension reduction (UMAP) representing the 6 major myeloid cell populations in which the 17 clusters analyzed using Flow-SOM were grouped depending on specific marker expression. (B) Frequency of monocytes (p=0.007, FDR=0.007) and dendritic cells (p=0.004, FDR=0.007) with respect to total myeloid cells among the LC and control cohorts. (C) Frequency of classical (p=n.s.), intermediate (p=n.s.) and non-classical (p=0.006, FDR=0.030) monocytes with respect to total monocytes among the LC and control cohorts. (D) Frequency of pDCs (p=0.043, FDR=0.071), cDCs (p=0.034, FDR=0.071) and CD123^−^ CD11c^−^ DCs (p=n.s.) with respect to total dendritic cells in the LC and control cohorts. (E) Frequency of Cluster 7 with respect to total cDCs among the LC and control cohorts (p=0.002, FDR=0.014). (F) Frequency of Cluster 11 with respect to total classical monocytes in the LC and control cohorts (p<0.001, FDR=0.001). (G) Mean fluorescence intensity (MFI) fold-change (FC) of CD169, CD38, CCR6, CCR7, CCR5, CXCR3 and CXCR5 in the LC and control cohorts in each of the different myeloid cell populations. * p < 0.05, FDR < 0.05. The frequency of each cluster was compared between the LC and control cohorts using linear regression models adjusted for SARS-CoV-2 antigen exposure, sex and age as covariates. The MFI of the different markers was compared between the LC and control cohorts using linear regression models adjusted for SARS-CoV-2 antigen exposure, sex and age as covariates, and a model-based FC was estimated between groups. LC n=98; controls n=18 in all analyses. Each dot represents an individual, and median and IQR values are indicated. p-values < 0.05 were considered statistically significant. p-values were adjusted for multiple testing using FDR.

### Quantification of SARS-CoV-2–specific IgG and IgA

To determine SARS-CoV-2–specific IgG and IgA titers, MaxiSorp ELISA plates were coated with anti-His tag monoclonal antibody (ThermoFisher, MA1-21315) at 2 μg/ml and incubated overnight at 4°C. The following day, plates were washed and blocked with 1% BSA in 1x PBS for two hours at room temperature. A half plate was then incubated with spike S2 subunit (S2), spike region binding domain (RBD) or nucleocapsid (N) (wild-type [Wuhan] strain; SinoBiological) at 1 μg/ml overnight at 4°C. The other half plate received antigen-free blocking buffer to determine sample background. Heat-inactivated (incubated at 56°C for 30 minutes) plasma samples were then added to the corresponding antigen-coated and antigen-free wells of the same plate and incubated for one hour at room temperature. Next, HRP-conjugated goat anti-human IgG or IgA (Jackson ImmunoResearch 109-036-098 and 109-035-011) was added to appropriate wells for 30 minutes at room temperature. Plates were then developed using o-phenylenediamine dihydrochloride (Sigma Aldrich, P8412-50TAB). The enzymatic reaction was stopped with 2N H2SO4. Absorbance was measured using the EnSight Multimode Plate Reader at 492 nm with noise correction at 620 nm. The specific signal was determined by subtracting the absorbance obtained in antigen-free wells from that of the wells coated with SARS-CoV-2 proteins. The results are expressed as arbitrary units per ml (AU/ml).

### Quantification of specific SARS-CoV-2 neutralizing capacity

Neutralization assays were performed in duplicate as previously described^15^. Briefly, 200 TCID50 of pseudovirus was preincubated in Nunc 96-well cell culture plates (Thermo Fisher Scientific), with three-fold serial dilutions (1/60–1/14,580) of heat-inactivated (incubated at 56°C for 30 minutes) plasma samples for 1 hour at 37°C. Then, 1×10^4^ HEK293T/hACE2 cells treated with DEAE-Dextran (Sigma-Aldrich) was added. The results were read after 48 hours using the EnSight Multimode Plate Reader and BriteLite Plus Luciferase reagent (PerkinElmer, USA). The values were normalized, and the ID50 (reciprocal dilution inhibiting 50% of the infection) was calculated by plotting and fitting all duplicate neutralization values and the log of the plasma samples dilution to a 4- parameter equation in Prism 9.0.2 (GraphPad Software, USA). This assay has previously been validated with a replicative viral inhibition assay^16^.

### Statistical analysis

Cell frequency, antibody levels and neutralizing capacity in each group were compared between the LC and control cohorts using linear regression models adjusted for SARS-CoV-2 antigen exposure, sex and age as covariates. For each cluster, the suitability of data transformation (logarithmic or square root) was assessed to approximate a normal distribution. The likelihood ratio test was used to compare nested models with and without the group factor. The same type of regression models, including covariate adjustment, was used to analyze differential expression of CD169, CD38, CCR6, CCR7, CCR5, CXCR3, CXCR5, PD1, CD25, CD95, CD8, HLADR, CD57, CD127 and CD1c, and model-based fold changes were estimated between groups. p-values <0.05 were considered statistically significant. In all analyses, p-values were adjusted for multiple testing using the false discovery rate.

When the LC and control groups were matched by time from last exposure, antibody levels and neutralizing capacity were compared using the Mann-Whitney test (non-parametric test); Spearman correlations were used for the correlation analysis.

In order to identify the markers with the highest discriminative capacity between LC and controls, a classification model based on random forest (RF) analysis was used, incorporating all the results for population frequency, differential expression analyses and antibody levels. The RF model was fitted with 5000 classification trees and tuned to account for class imbalance between the two cohorts. The number of markers selected was determined through multiple selection strategies based on their importance scores, ranging from more to less conservative. The RF model was refitted using only the selected markers in each case, and the resulting models were compared based on their out-of-bag estimate of error rate. Finally, a heatmap combined with hierarchical clustering was used to visually summarize patterns of marker expression across samples and differences between cohorts.

A survival analysis was conducted to analyze the potential relationship between the classification made by the RF model within the LC cohort (based on immunological markers) and the clinical severity of the disease. This analysis aimed to determine whether LC samples misclassified by the RF model were associated with different clinical conditions. It focused on the time from the LC diagnosis to medical discharge, estimating survival curves using the Kaplan-Meier method and plotting the cumulative recovery curve accordingly.

The analyses were performed using R version 4.4.1 and GraphPad Prism version 10.2.2 for Windows (GraphPad Software, Boston, Massachusetts USA, www.graphpad.com).

Only p-values <0.05 were considered statistically significant.

## RESULTS

### Characteristics of the pediaCOVID cohort

The pediaCOVID cohort^6,17^ is composed of two groups: the LC group, comprising 99 CYP diagnosed with LC; and a control group, comprising 18 CYP without LC or any other inflammatory disease or acute infection. The clinical characteristics of both groups are described in Table 1.

**Table 1:** Clinical characteristics of the pediaCOVID cohort.

|  | Long COVID cohort<br>n = 99 | Control cohort<br>n = 18 |
| --- | --- | --- |
| Age at sample collection (years), median (IQR) | 13.9 (12.3-15.0) | 13.9 (10.2-16.6) |
| Female sex, n (%) | 63 (63.6) | 7 (38.9) |
| One exposure to SARS-CoV-2 antigens, n (%)<sup>a</sup> | 83 (83.8) | 15 (83.3) |
| Two exposures to SARS-CoV-2 antigens, n (%)<sup>b</sup> | 16 (16.2) | 3 (16.7) |
| Time since acute SARS-CoV-2 infection (weeks), median (IQR)<sup>c</sup> | 23.5 (16.3-39.2) | 41.4 (13.6-60.4) |
| Time since last SARS-CoV-2 antigen exposure (weeks), median (IQR)<sup>d</sup> | 20.3 (13.9-34.7) | 11.6 (8.1-17.3) |
| Number of persistent symptoms after acute COVID-19, median (IQR) | 12 (8-17) | - |
| **General symptoms** |  |  |
| Fatigue, n (%) | 97 (98.0%) | - |
| Skin rashes, n (%) | 22 (22.2%) | - |
| Fever, n (%) | 10 (10.1%) | - |
| COVID toes, n (%) | 1 (1.0%) | - |
| **Neurological and neurocognitive impairment symptoms** |  |  |
| Headache, n (%) | 75 (75.8%) | - |
| Attention and planning disturbances, n (%) | 51 (51.5%) | - |
| Lack of concentration, n (%) | 51 (51.5%) | - |
| Brain fog, n (%) | 47 (47.5%) | - |
| Insomnia, n (%) | 45 (45.5%) | - |
| Memory disturbances, n (%) | 39 (39.4%) | - |
| Sonophobia, n (%) | 37 (37.4%) | - |
| Photophobia, n (%) | 35 (35.4%) | - |
| Anosmia, n (%) | 34 (34.3%) | - |
| Paresthesia, n (%) | 32 (32.3%) | - |
| Ageusia, n (%) | 30 (30.3%) | - |
| Vertigo, n (%) | 24 (24.2%) | - |
| Dizziness, n (%) | 7 (7.1%) | - |
| Dysphonia, n (%) | 1 (1.0%) | - |
| **Respiratory symptoms** |  |  |
| Dyspnea, n (%) | 63 (63.6%) | - |
| Cough, n (%) | 24 (24.2%) | - |
| Nasal congestion, n (%) | 2 (2.0%) | - |
| **Cardiovascular symptoms** |  |  |
| Orthostatic hypotension, n (%) | 46 (46.5%) | - |
| Thoracic pain, n (%) | 39 (39.4%) | - |
| Tachycardia, n (%) | 38 (38.4%) | - |
| Palpitations, n (%) | 31 (31.3%) | - |
| **Gastrointestinal symptoms** |  |  |
| Abdominal pain, n (%) | 39 (39.4%) | - |
| Diarrhea, n (%) | 20 (20.2%) | - |
| Vomiting and/or nausea, n (%) | 15 (15.2%) | - |
| Dysphagia, n (%) | 10 (10.1%) | - |
| Constipation, n (%) | 3 (3.0%) | - |
| **Musculoskeletal symptoms** |  |  |
| Muscle weakness, n (%) | 64 (64.6%) | - |
| Myalgia/arthralgia, n (%) | 56 (56.6%) | - |
| Decreased strength in upper extremities, n (%) | 32 (32.3%) | - |
| **Eating disorders** | 35 (35.4%) | - |
| **Menstrual disorders, n (%)** | 17 (17.2%) | - |
| <sup>a</sup>SARS-CoV-2–infected or –vaccinated individuals |  |  |
| <sup>b</sup>SARS-CoV-2–infected and –vaccinated individuals |  |  |
| <sup>c</sup>Data only from individuals with date of SARS-CoV-2 infection |  |  |
| <sup>d</sup>Antigen exposure: Infection or vaccination |  |  |

### Myeloid cells are dysregulated in CYP with LC

In total, 80 million cells were analyzed using spectral flow cytometry. Of these, 15% were myeloid cells, 12% were NK cells, 65% were T cells and 8% were B cells (Supp.Table.6).

Seventeen different clusters were identified in the myeloid cell population using the following markers: CD1c, CD11b, CD16, CD11c, HLADR, CD95, CD14 and CD123 (Supp.Fig.2, Supp.Table.2). Each cluster was annotated as a monocyte or dendritic cell (DC) cluster depending on the expression of various markers (Supp.Table.6). Monocyte clusters were classified as classical, intermediate or non-classical, while DC clusters were classified as conventional (cDCs), plasmacytoid (pDCs) or CD11c^−^ CD123^−^ DCs (Figure.1a, Supp.Table.6).

**Figure 2:**
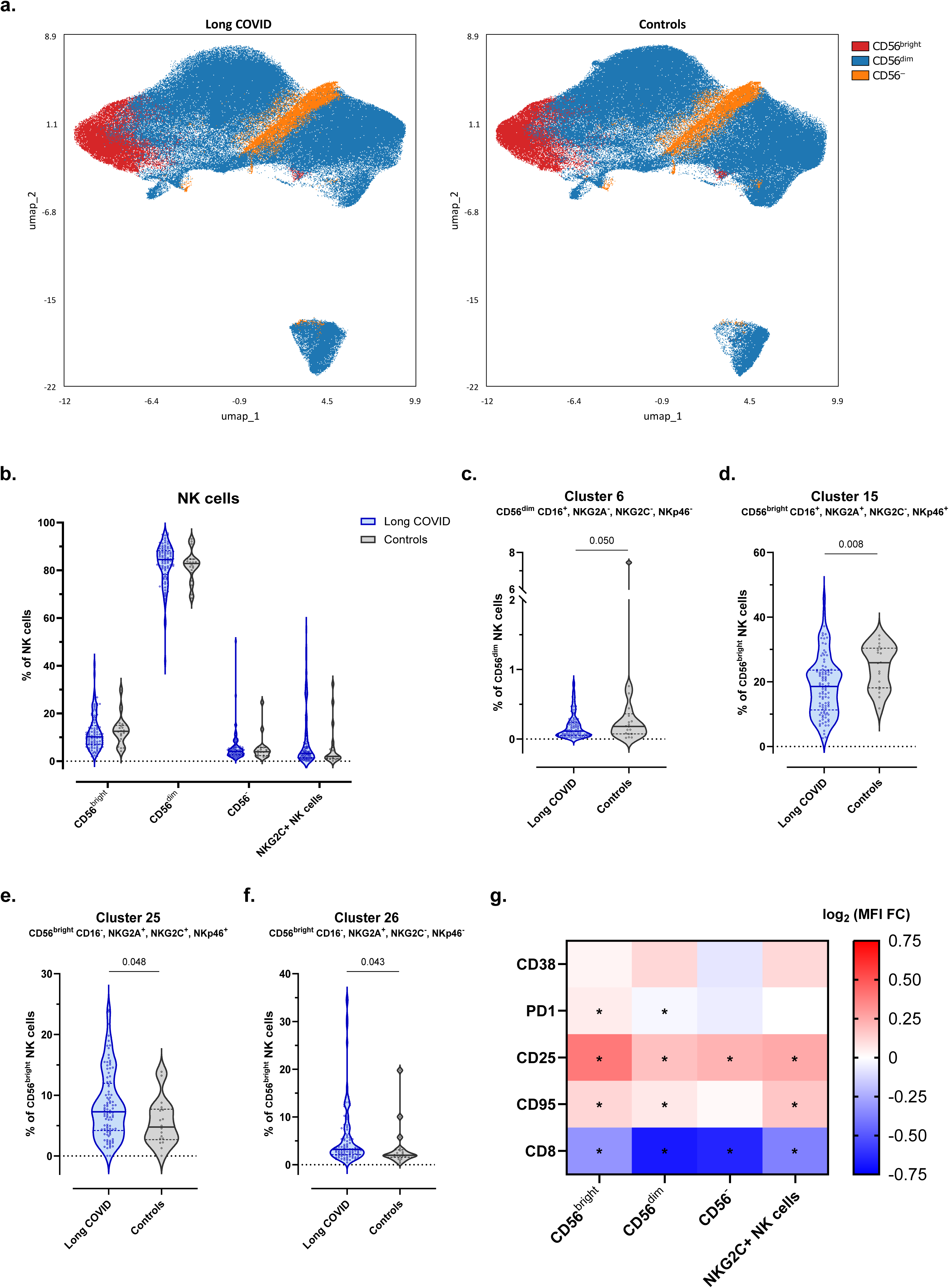
NK-cell compartment in CYP with and without LC. (A) Uniform manifold approximation and projection for dimension reduction (UMAP) representing the 4 major NK-cell populations into which the 33 clusters analyzed using Flow-SOM were grouped depending on specific marker expression. (B) Frequency of CD56^bright^, CD56^dim^, CD56^−^ and NKG2C^+^ NK cells with respect to total NK cells (all p=n.s.). (C) Frequency of Cluster 6 with respect to total CD56^dim^ NK cells in the LC and control cohorts (p=0.050, FDR=0.427). (D) Frequency of Cluster 15 with respect to total CD56^bright^ NK cells in the LC and control cohorts (p=0.008, FDR=0.306). (E) Frequency of Cluster 25 with respect total CD56^bright^ NK cells in the LC and control cohorts (p=0.048, FDR=0.427). (F) Frequency of Cluster 26 with respect to total CD56^bright^ NK cells in the LC and control cohorts (p=0.043, FDR=0.427). (G) Mean fluorescence intensity (MFI) fold-change (FC) of CD38, PD1, CD25, CD95 and CD8 between the LC and control cohorts in each of the different NK cell populations. * p < 0.05, FDR < 0.05. The frequency of each cluster was compared between the LC and control cohorts using linear regression models adjusted for SARS-CoV-2 antigen exposure, sex and age as covariates. The MFI of the different markers was compared between the LC and control cohorts using linear regression models adjusted for SARS-CoV-2 antigen exposure, sex and age as covariates, and a model-based FC was estimated between the groups. LC n=98; controls n=18 in all analyses. Each dot represents an individual, and median and IQR values are indicated. p- values < 0.05 were considered statistically significant. p-values were adjusted for multiple testing using FDR.

Pediatric LC was associated with decreased frequency of monocytes and increased frequency of DCs (Figure.1b). Sub-analysis of the three different monocyte populations revealed increased frequency of non-classical monocytes in CYP with LC (Figure.1c). The DC compartment was also dysregulated in pediatric LC, with higher proportions of pDCs and decreased cDCs (Figure.1d). Among the different clusters analyzed, cluster 7 (CD14^+^, CD11c^+^, CD1c^+^) was decreased in pediatric LC, while cluster 11 (CD14^+^, CD16^−^, CD123^+^) was increased (Figure.1e-f).

Finally, we analyzed the differential expression of various markers (CD169, CD38, CCR7, CCR6, CCR5, CXCR3 and CXCR5) between CYP with and without LC (Figure.1g). Pediatric LC was associated with decreased expression of both CCR7 and CCR6 in DCs and monocyte populations. Decreased expression of CXCR5, CD169 and CD38 was associated with pediatric LC, albeit only in some DC and monocyte populations.

Pediatric LC was associated with a dysregulated myeloid cell compartment marked by a lower frequency of monocytes that were enriched in the non-classical subset and an increase in DCs with an overrepresentation of pDCs. Both monocytes and DCs showed lower expression of CCR6 and CCR7 than the control group.

### Pediatric LC is associated with increased NK-cell activation

Thirty-three different clusters were identified in the NK-cell population using the following markers: CD56, CD16, CD57, NKG2A, NKG2C, NKp46, CCR5 and CXCR3 (Supp.Fig.2, Supp.Table.3). Each cluster was then annotated as CD56^bright^, CD56^dim^, CD56^−^ or NKG2C+ memory NK cells (Figure.2a, Supp.Table.6).

No significant differences were recorded for any of the major NK-cell populations between CYP with and without LC (Figure.2b). Pediatric LC was associated with decreased levels of cluster 6 (CD56^dim^ NK-cell CD16^+^, NKG2A^−^, NKG2C^−^, NKp46^−^) (Figure.2c) and cluster 15 (CD56^bright^ NK-cell CD16^+^, NKG2A^+^, NKG2C^−^, NKp46^+^) (Figure.2d) and with increased levels of cluster 25 (CD56^bright^ NK-cell CD16^−^, NKG2A^+^, NKG2C^+^, NKp46^+^) (Figure.2e) and cluster 26 (CD56^bright^ NK-cell CD16^−^, NKG2A^+^, NKG2C^−^, NKp46^−^) (Figure.2f).

The differential expression of CD28, PD1, CD25, CD95 and CD8 was analyzed in the different NK-cell populations by comparing the level of each marker in CYP with and without LC (Figure.2g). Pediatric LC was associated with higher expression of CD25 and decreased expression of CD8 in all NK-cell populations. PD1 and CD95 were also increased in pediatric LC but not in all the NK-cell populations.

The NK-cell compartment in pediatric LC shifted from cytotoxic CD16^+^ subsets toward CD16^−^ NK cells, suggesting a dysregulated profile with increased activation but impaired maturation and function.

### T cells are more frequently activated and exhausted but dysregulated in pediatric LC

Thirty-seven different clusters were identified in the T-cell population using the following markers: TCRVd2, CD4, CD8, CD45RA, CCR7, CD95, CD27, CD28, CCR6, CXCR3, CD25 and CD127 (Supp.Fig.2, Supp.Table.4). Each cluster was annotated as a γδ T cell, CD4 T cell, CD8 T cell, double-positive T cell or double-negative T cell (Figure.3a). Then, CD4 and CD8 T-cell clusters were assigned to the different memory, naïve and regulatory subpopulations through expression of the markers CD45RA, CCR7, CD27, CD28, CD95, CD25 and CD127 (Supp.Table.6).

Pediatric LC was associated with an increase in CD4 T cells and a trend towards a decrease in CD8 T cells (Figure.3b). Moreover, the CD4:CD8 ratio was higher in CYP with LC than in controls, although it did not reach statistical significance (data not shown).

In the CD4 T-cell subpopulations, pediatric LC was associated with a decrease in central memory CD4 T cells (CD4 TCM) and an increase in regulatory CD4 T cells (CD4 Treg) (Figure.3c,e). Moreover, the two clusters that formed the CD4 Treg population were dysregulated in pediatric LC, with an expansion of cluster 3 (naïve CD4 Treg CD45RA^+^, CD95^−^) and a decreased proportion of cluster 25 (memory/activated CD4 Treg CD45RA^−^, CD95^+^) (Figure.3e).

In the CD8 T-cell compartment, pediatric LC was associated with a decrease in stem cell– like memory CD8 T cells (CD8 TSCM) and increased central memory CD8 T cells (CD8 TCM) (Figure.3d,f). Among all the clusters, the two clusters that formed the CD8 TCM population were dysregulated in CYP with LC, with increased proportions of cluster 16 (CD8 TCM CXCR3^+^, CCR6^+^) and decreased proportions of cluster 23 (CD8 TCM CXCR3^−^, CCR6^−^) (Figure.3f).

Finally, the differential expression of CD25, HLADR, CD38, PD1 and CD57 was analyzed in the various T-cell populations between CYP with and without LC (Figure.3g). Pediatric LC was associated with higher expression of activation (HLA-DR, CD38, CD25) and senescence/exhaustion (CD57, PD-1) markers, particularly in the CD4 TCM, CD8 TCM and γδ T-cell populations.

Pediatric LC was associated with dysregulation of the T-cell compartment driven by decreased CD4 TCM and CD8 TSCM and increased CD4 Treg and CD8 TCM. Moreover, CYP with LC had higher levels o f T-cell activation characterized by a higher expression of CD25, HLADR, CD38, PD1 and CD57.

### Impaired humoral response against SARS-CoV-2 in pediatric LC

Thirty-one clusters were identified in the B-cell population using the following markers: IgD, IgM, IgA, CD27, CD21, CD38, CD24, CD11c, CCR7, CCR6, CXCR5 and CXCR3 (Supp.Fig.2, Supp.Table.5). Each cluster was annotated as transitional B cells, naïve B cells, marginal zone (MZ)–like B cells, IgM-only B cells, IgD-only B cells, switched memory B cells and plasmablasts (Figure.4a-b, Supp.Table.6).

Pediatric LC was associated with a higher frequency of switched memory and IgD-only B cells (Figure.4c). Specifically, we observed an increase in the frequency of switched memory B cells from cluster 15 (IgA^−^CD27^−^CD21^−^), which lacked the expression of CD27 and CD21, at the expense of the CD27^+^ switched memory B-cell subsets (cluster 17 [switched memory B cells CD27^+^CD21^+^CD24^hi^] and cluster 19 [switched memory B cells CD27^+^CD21^+^CD24^−^]) (Figure.4d). While no differences in the size of the MZ-like B-cell compartment were detected, the frequency of MZ-like B cells expressing CXCR3 (cluster 25) was lower in pediatric LC (Figure.4d).

When the expression of CD127, CD1c, CD25, PD1 and CD95 was analyzed in CYP with and without LC (Figure.4e), pediatric LC was associated with decreased expression of CD1c in all B-cell populations. Interestingly, PD1 expression was increased in all B-cell populations except plasmablasts, where it decreased. Similarly, CD25 was increased in transitional, naïve, IgD-only and switched memory B cells but decreased in plasmablasts (Figure.4e).

These results prompted us to check specific antibody levels against SARS-CoV-2 in CYP with and without LC. For that purpose, we used ELISA to quantify plasma levels of IgG and IgA antibodies against SARS-CoV-2 antigens (spike S2, RBD and N). Pediatric LC was associated with lower levels of anti-RBD IgG and IgA antibodies (Figure.4f,i). However, we observed no significant differences in anti-S2 IgG and IgA and anti-N IgG antibody levels between the two cohorts (Figure.4f). As expected, CYP with two exposures (vaccinated after acute infection) had significantly higher levels of anti-S2 and anti-RBD antibodies than CYP with one exposure (either infection or vaccination) (Supp.Table.7).

To determine whether neutralizing capacity was impaired owing to lower anti-RBD response, we quantified the level of neutralization using a luciferase-reporter lentiviral pseudovirus assay. Pediatric LC was associated with lower levels of neutralizing antibodies against the D614G variant (Figure.4l). However, these increased in CYP with LC after two exposures, as expected (Supp.Table.7). Moreover, in CYP with LC, levels of neutralizing antibodies correlated positively with levels of both anti-RBD and anti-S2 IgG and IgA antibodies (Figure.4m). For the control cohort, only anti-RBD IgG levels correlated with neutralizing capacity (Figure.4m).

A critical factor to consider when analyzing antibody levels is the time since the most recent antigen exposure (either through vaccination or natural infection), as antibody levels typically diminish over time. Sample collection in the control cohort was closer to the most recent antigen exposure (median 11.6 weeks; IQR [8.1-17.3]) than in the LC cohort (median 20.3 weeks; IQR [13.9-34.7]) (Suppl.Fig.3a). However, despite these differences, no correlation was observed between anti-RBD–specific IgG/IgA levels and the time since the last exposure in either the LC or the control cohort with a single-exposure event (Suppl.Fig.3.b). Antibody levels and neutralizing capacity in CYP with LC remained significantly lower when the sample time point of CYP with LC was restricted to fewer than 17 weeks since the last antigen exposure to fit within the IQR of the control cohort, demonstrating that time from the most recent antigen exposure did not affect the differences observed between the LC and the control cohort (Suppl.Fig.3c-e).

**Figure 3:**
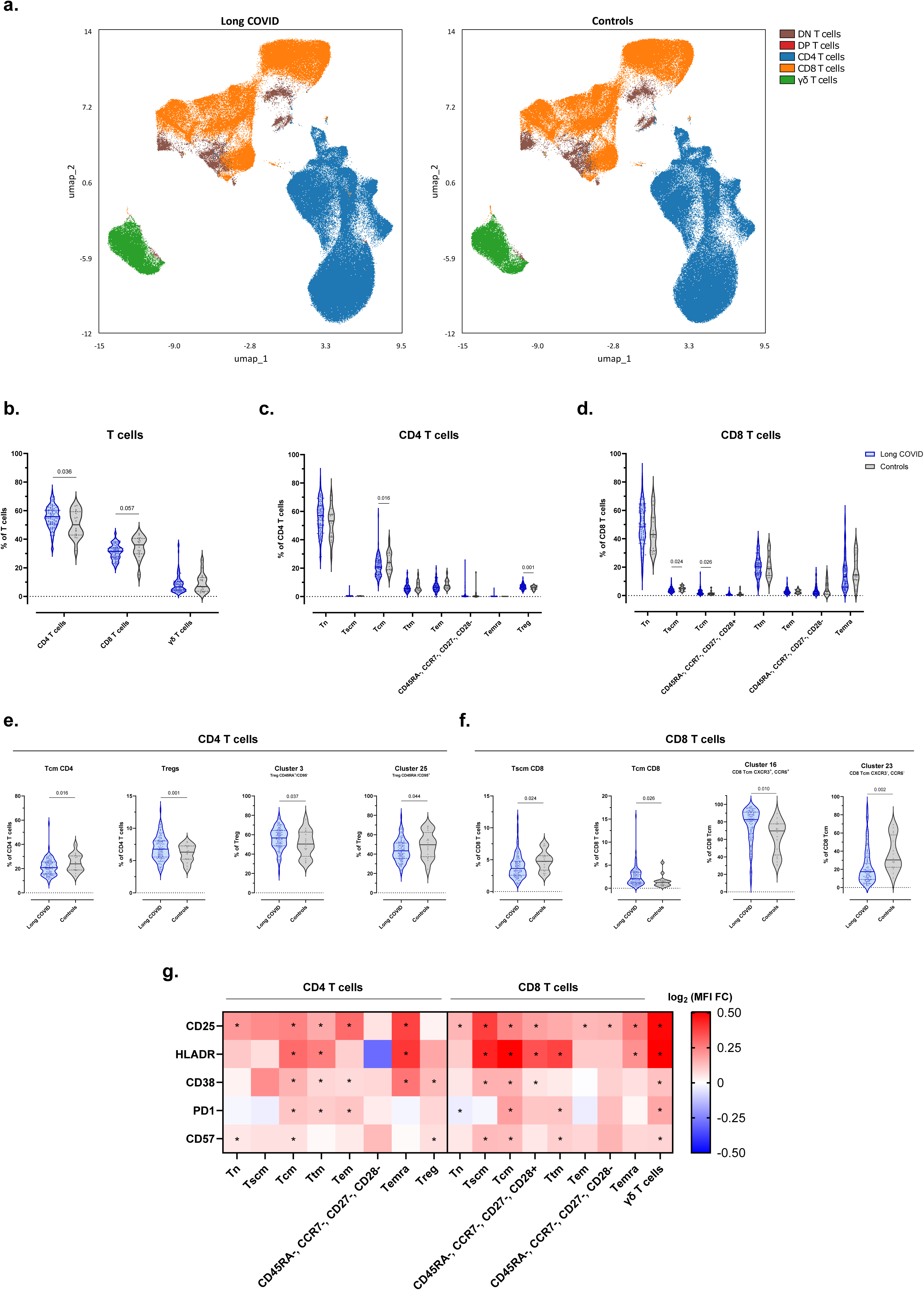
T-cell compartment in CYP with and without LC. (A) Uniform manifold approximation and projection for dimension reduction (UMAP) representing the 5 major T-cell populations into which the 37 clusters analyzed using Flow-SOM were grouped depending on specific marker expression. (B) Frequency of CD4 (p=0.036, FDR=0.087), CD8 (p=0.057, FDR=0.087) and γδ T cells (p=n.s.) with respect to total T cells in the LC and control cohorts. (C) Frequency of CD4 subpopulations with respect to total CD4 T cells in the LC and control cohorts (CD4 Tcm p=0.016, FDR=0.118; CD4 Treg p=0.029, FDR=0.118; Other p=n.s.). (D) Frequency of CD8 subpopulations with respect to total CD8 T cells in the LC and control cohorts (CD8 Tscm p=0.024, FDR=0.118; CD8 Tcm p=0.026, FDR=0.118; Other p=n.s.). (E) From left to right; Frequency of Tcm CD4 (p=0.016, FDR=0.118) with respect to total CD4 T cells, frequency of Treg (p=0.029, FDR=0.118) with respect to total CD4 T cells, frequency of Cluster 3 (p=0.037, FDR=0.274) with respect to total Treg and frequency of Cluster 25 (p=0.044, FDR=0.274) with respect to total Treg in the LC and control cohorts. (F) From left to right; Frequency of Tscm CD8 (p=0.024, FDR=0.118) with respect to total CD8 T cells, frequency of Tcm CD8 (p=0.026, FDR=0.118) with respect to total CD4 T cells, frequency of Cluster 16 (p=0.010, FDR=0.126) with respect to total Tcm CD8 and frequency of Cluster 23 (p=0.002, FDR=0.058) with respect to total Tcm CD8 among the LC and control cohorts. (G) Mean fluorescence intensity (MFI) fold-change (FC) of CD25, HLADR, PD1, CD38 and CD57 between the LC and control cohorts in each of the different T-cell populations. * p < 0.05, FDR < 0.05. The frequency of each cluster was compared between the LC and control cohorts using linear regression models adjusted for SARS-CoV-2 antigen exposure, sex and age as covariates. The MFI of the different markers was compared between the LC and control cohorts using linear regression models adjusted for the SARS-CoV-2 antigen exposure, sex and age as covariates, and a model-based FC was estimated between the groups. LC n=97; controls n=16 in all analyses. Each dot represents an individual, and median and IQR values are indicated. p-values < 0.05 were considered statistically significant. p-values were adjusted for multiple testing using FDR.

**Figure 4:**
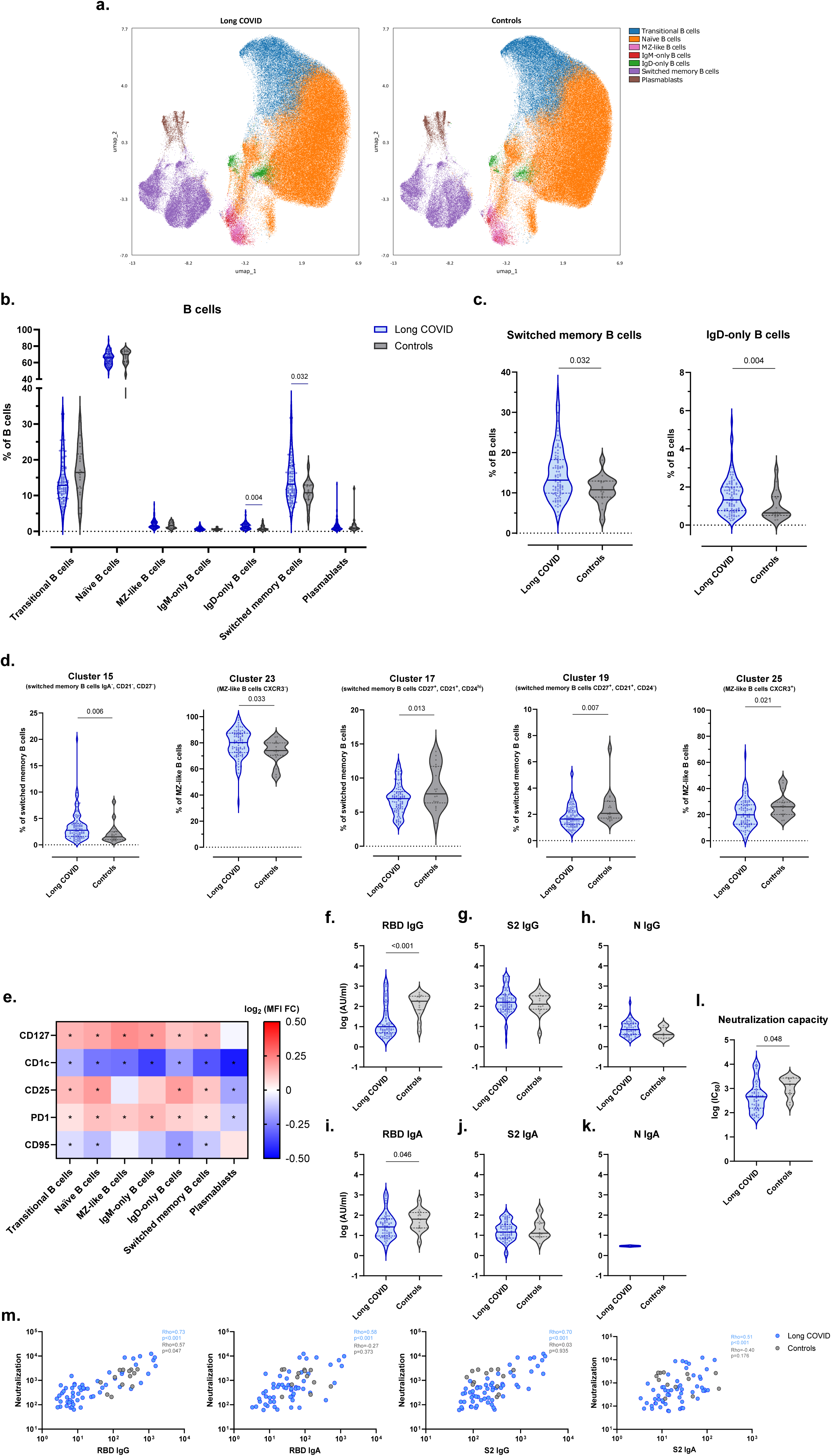
Humoral response in CYP with and without LC. (A) Uniform manifold approximation and projection for dimension reduction (UMAP) representing the 6 major B-cell populations in which the 31 clusters analyzed using Flow-SOM were grouped depending on specific marker expression. (B) Frequency of major B-cell populations with respect to total B cells among the LC and the control cohort (IgD-only B cells p=0.004, FDR=0.029; Switched memory B cells p=0.032, FDR=0.110; Other p=n.s.). (C) From left to right; Frequency of switched memory (p=0.032, FDR=0.110) and IgD-only B cells (p=0.004, FDR=0.029) with respect to total B cells in the LC and the control cohorts. (D) From left to right; Frequency of Cluster 15 (p=0.006, FDR=0.115) with respect to total switched memory B cells, frequency of Cluster 23 (p=0.033, FDR=0.202) with respect to total MZ-like B cells, frequency of Cluster 17 (p=0.013, FDR=0.127) with respect to total switched memory B cells, frequency of Cluster 19 (p=0.007, FDR=0.115) with respect to total switched memory B cells and frequency of Cluster 25 (p=0.021, FDR=0.162) with respect to total MZ-like B cells in the LC and control cohorts. (E) Mean fluorescence intensity (MFI) fold-change (FC) of CD127, CD1c, CD25, PD1 and CD95 between LC and control cohorts in each of the different B-cell populations. * p < 0.05, FDR < 0.05. (F) Anti-RBD IgG antibody levels (p<0.001) in responders in log (AU/ml). (G) Anti-S2 IgG antibody levels (p=n.s.) in responders in log (AU/ml). (H) anti-N IgG antibody levels (p=n.s.) in responders in log (AU/ml). (I) Anti-RBD IgA antibody levels (p=0.046) in responders in log (AU/ml). (J) Anti-S2 IgA antibody levels (p=n.s.) in responders in log (AU/ml). (K) Anti-N IgA antibody levels (p=n.s.) in responders in log (AU/ml). (L) SARS-CoV-2 neutralizing capacity (p=0.048) of responders in log/half-maximal inhibitory concentration [IC50]) in the LC and control cohorts. (M) From left to right; Correlation between neutralizing capacity (IC50) and anti-RBD IgG antibodies (AU/ml) (LC cohort, anti-RBD IgG/Neutralizing capacity, p < 0.001, Rho = 0.73; control cohort, anti-RBD IgG/Neutralizing capacity, p = 0.047, Rho = 0.57), correlation between neutralizing capacity (IC50) and anti-RBD IgA antibodies (AU/ml) (LC cohort, anti-RBD IgA/Neutralizing capacity, p < 0.001, Rho = 0.58; control cohort, anti-RBD IgA/Neutralizing capacity, p=n.s., Rho = -0.27), correlation between neutralizing capacity (IC50) and anti-S2 IgG antibodies (AU/ml) (LC cohort, anti-S2 IgG/Neutralizing capacity, p < 0.001, Rho = 0.70; control cohort, anti-S2 IgG/Neutralizing capacity, p=n.s. Rho = 0.03), correlation between neutralizing capacity (IC50) and anti-S2 IgA antibodies (AU/ml) (LC cohort, anti-S2 IgA/Neutralizing capacity, p < 0.001, Rho = 0.51; control cohort, anti-S2 IgA/Neutralizing capacity, p=n.s., Rho = –0.40) in the LC and control cohort. The frequency of each cluster was compared between the LC and control cohorts using linear regression models adjusted for SARS-CoV-2 antigen exposure, sex and age as covariates. The MFI of the different markers was compared between the LC and control cohorts using linear regression models adjusted for SARS-CoV-2 antigen exposure, sex and age as covariates, and a model-based FC was estimated between groups. LC n=93; controls n=18 in the B-cell analysis. Antibody levels and neutralizing capacity were compared between the LC and control cohorts using linear regression models adjusted for SARS-CoV-2 antigen exposure, sex and age as covariates. Each dot represents an individual, and median and IQR values are indicated. p-values < 0.05 were considered statistically significant. p-values were adjusted for multiple testing using FDR.

Overall, pediatric LC was characterized by dysregulation of the B-cell compartment and an impaired anti-RBD humoral response associated with decreased neutralizing activity against SARS-CoV-2.

### Decreased CCR6 expression in monocytes distinguishes LC patients from controls

To identify immunophenotypic features distinguishing LC from controls, we used random forest (RF)–based classification. The RF model achieved an overall accuracy of 79.3% in differentiating LC cases from controls (Supp.Fig.5). Among all the parameters analyzed, the nine most influential were the following: CCR6 expression in classical, total and intermediate monocytes; CXCR5 expression in CD4 TEM and cDCs; CD8 expression in CD56^−^ NK cells; CD1c expression in MZ-like and IgM-only B cells; and CCR6 expression in CD56^bright^ NK cells (Figure.5a). Notably, CCR6 expression on classical, total and intermediate monocytes were the most significant factors for distinguishing LC from control individuals. Moreover, we observed that individuals diagnosed with LC who were misclassified as controls by the RF model exhibited a higher rate of recovery than those classified correctly (96.0% vs 71%, p=0.012) (Figure.5b). However, there were no significant differences in the time from diagnosis to recovery between the two groups (Figure.5c).

**Figure 5:**
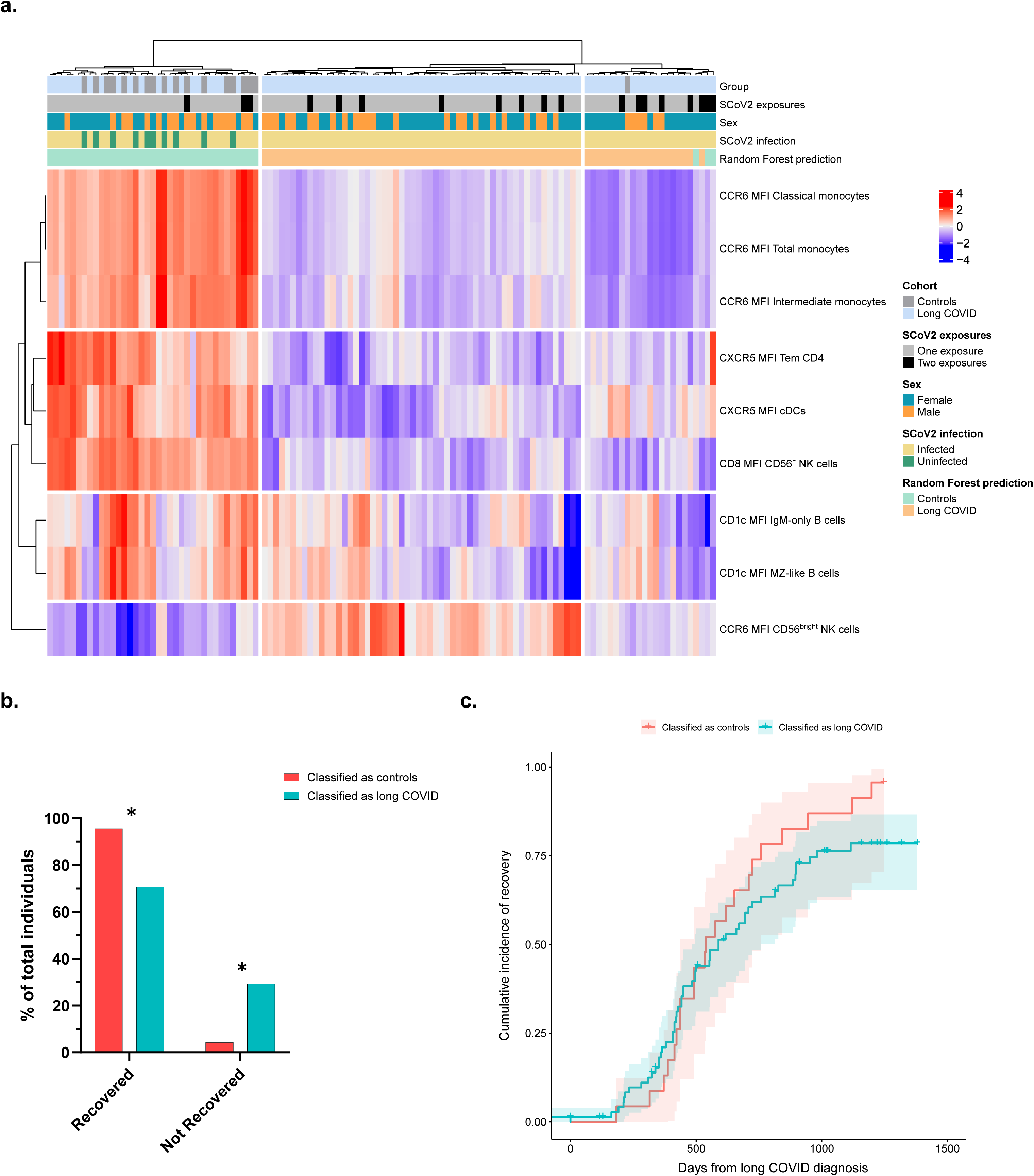
Random forest model classifies long COVID patients and controls with 79.2% accuracy. (A) Heatmap representing the nine most important features that distinguish long COVID patients from controls in the random forest model (from most important CCR6 MFI classical monocytes to least important CCR6 MFI CD56^bright^ NK cells) (LC n=98; controls n=18). (B) Percentage of recovered individuals in long COVID patients who were classified as controls or LC by the random forest model (classified as controls n=23; classified as LC n=75; * p=0.012, Fisher’s exact test). (C) Kaplan-Meier curves showing the proportion of unrecovered long COVID patients over time in individuals classified as controls or LC by the random forest model (classified as controls n=23; classified as LC n=75; p=n.s.).

## DISCUSSION

Elucidating the pathophysiology of LC and identifying reliable biomarkers remain challenging in both children and adults. In this study, the comprehensive analyses of PBMCs and plasma from CYP with and without LC revealed significant immunological differences.

Immunophenotyping of the major immune cell populations (myeloid, NK, T and B cells) revealed significant differences between CYP with and without LC. Notably, pediatric LC was associated with an increased proportion of DCs relative to monocytes. Although the overall proportion of monocytes was reduced, there was a specific increase in non-classical monocytes among CYP with LC. Non-classical monocytes are widely implicated in chronic inflammation and autoimmunity and have previously been reported to be elevated in adults with LC^18,19^. Moreover, the proportion of pDCs and cDCs was altered in pediatric LC, with higher levels of pDCs and lower levels of cDCs than in controls. pDCs are the primary producers of type I interferon responses, and an aberrantly high or delayed pDC response in COVID-19 has been linked to tissue damage^20^. In contrast, cDCs function primarily in antigen presentation to initiate adaptive immune responses^21^. Importantly, cluster 7 (cDC, CD14^+^, CD1c^+^), which phenotypically resembles DC3 subset^22^, was significantly reduced in pediatric LC. DC3 has been shown to stimulate T cells as effectively as other cDC subsets, specifically polarizing CD8 T cells to CD8 resident memory (CD103^+^)^22^. Interestingly, expression of CCR7 and CCR6 was reduced in all major populations of DCs and monocytes; expression of CXCR5 and CD169 was reduced in most major populations. Through its interaction with CCL20, CCR6 plays a critical role in the recruitment of DCs and monocytes to inflamed mucosal tissues^23^. Conversely, CCR7 is upregulated during DC maturation, enabling its migration to lymph nodes upon recognition of the ligands CCL19 and 21^24^. Interestingly, during acute SARS-CoV-2 infection, DCs display functional impairment, which is characterized by diminished expression of maturation markers such as HLADR, CD86 and CCR7 following stimulation^25^. Diminished CCR6 expression could impair the ability of antigen-presenting cells (APCs) to migrate to inflamed tissues, thereby disrupting antigen processing and resulting in reduced maturation. This, in turn, may lower CCR7 expression, further impairing migration of APCs to secondary lymph organs and ultimately weakening the induction of robust T- and B-cell responses. Finally, reduced CCR6 levels may also hinder migration of APCs to inflamed mucosal tissues (e.g. gut), resulting in diminished local immune responses. These findings are consistent with those of previous studies reporting persistence of SARS-CoV-2 in the gut and other tissues^12,26^.

No significant differences were observed in the major NK-cell populations between CYP with and without LC (CD56^bright^, CD56^dim^, CD56^−^). However, among specific NK cell clusters, four were found to be dysregulated in pediatric LC. Of note, the clusters that decreased in pediatric LC were CD16^+^, whereas those that increased did not express CD16. CD16 (FcγRIII) is a key receptor mediating antibody-dependent cellular cytotoxicity, which is critical for cytotoxic function in NK cells^27^. Finally, pediatric LC was associated with increased CD25 expression and reduced CD8 expression across all NK-cell subsets. CD25 is upregulated upon NK-cell activation and has been shown to enhance cytotoxic activity^28^. In contrast, CD8 expression was lower in all NK-cell subsets. Although CD8 is primarily expressed by CD8 T cells, it is also present on some NK cells^29^. Moreover, CD8^+^ NK cells have been reported to have greater functional activity than CD8^−^ NK cells in people living with HIV-1^30^.

Pediatric LC was associated with an increased frequency of CD4 T cells. Moreover, CYP with LC had elevated levels of Treg alongside reduced CD4 TCM. Interestingly, these features have previously been described in adults with LC^19,31^. Analysis of the Treg population revealed pediatric LC to be associated with two distinct clusters, namely, increased levels of CD45RA^+^ Treg and decreased levels of CD45RA^−^ Treg. CD45RA^+^ Treg maintain a more naïve phenotype, whereas CD45RA^−^ adopt a more memory-like phenotype with enhanced proliferative capacity^32^. Notably, 90-95% of Foxp3^+^ cells are CD45RA^−^, indicating that they have already undergone activation and proliferation^33^. In the CD8^+^ T-cell compartment, pediatric LC was characterized by a reduction in CD8 TSCM and increased CD8 TCM compared to controls. CD8 TCM are critical for long-term immune protection owing to their high proliferative potential, persistence in blood, and ability to generate effector and effector memory T cells upon re-exposure to antigen^34,35^. Recent evidence also suggests that TCM play a more important role in managing systemic infections than TEM, as they are centrally involved in secondary lymphoid tissues and possess greater proliferative capacity^36^. Additionally, CYP with LC showed increased levels of CD8 TCM expressing CXCR3 and CCR6, while CD8 TCM lacking these markers were less common than in the control cohort. This observation is noteworthy, as elevated CXCR3 expression on CD8 TCM has also been reported in people living with HIV-1, suggesting a shared trait of chronic viral infections^37^. Finally, pediatric LC was associated with heightened activation and senescence/exhaustion in the CD4, CD8 and γδ T-cell compartments, as evidenced by more marked expression of CD25, HLADR, CD38, PD1 and CD57 than controls. Interestingly, CD4 TCM, CD8 TCM and γδ T cells all highly expressed these five markers in CYP with LC, indicating pronounced activation and exhaustion within these specific subsets.

B-cell immunophenotyping revealed that CYP with LC exhibited higher frequencies of IgD-only B cells and total switched memory B cells than the control cohort. A previous study reported no significant differences in the B-cell compartment between children with LC and recovered individuals^38^. However, the authors analyzed a more limited panel of markers to characterize B-cell subsets, in contrast to the broader profiling used in our analysis. Interestingly, IgD expressed by IgD-only B cells has been shown to be enriched in autoreactivity^39^. Nevertheless, the functional role of these specific B-cell subsets remains unclear, and further studies are warranted to determine their relevance in LC. Despite the overall increase in total switched memory B cells in pediatric LC, we identified a dysregulated IgG^+^ memory B-cell compartment. This was characterized by elevated levels of IgA^−^CD21^−^CD27^−^ (cluster 15) and reduced levels of IgA^−^ CD21^+^CD27^+^CD24^hi^ (cluster 17) and IgA^-^CD21^+^CD27^+^CD24^−^ (cluster 19) switched memory B cells. Cluster 15 cells are known to express inhibitory receptors such as FcRL4 and FcRL5 and display an “exhausted-like” phenotype, commonly observed in chronic viral infections such as HIV-1^40^. Although FcRL4 expression was not directly measured in our study, FcRL4^+^ B cells are less likely to differentiate into antibody-secreting plasma cells^41^. Notably, CD21^−^CD27^−^ memory B cells can arise from extrafollicular humoral responses or from primary germinal center reactions^42^, suggesting that CYP with LC may undergo impaired germinal center maturation. Conversely, the CD21^+^CD27^+^ subsets (clusters 17 and 19) are typically associated with higher proliferative capacity, increased somatic hypermutation, and robust antibody production upon re-exposure to antigen^43^. The observed expansion of CD21^−^CD27^−^ and contraction of CD21^+^CD27^+^ memory B cells is a hallmark of acute COVID-19, although it is typically resolved in recovered individuals^44^. This persistence in pediatric LC suggests sustained impairment of humoral immune function.

Furthermore, relative to controls, CYP with LC had an increased frequency of MZ-like B cells lacking CXCR3 expression and a decreased frequency of CXCR3^+^ MZ-like B cells. However, total MZ-like B-cell frequency remained unchanged between the groups. The primary distinction between these two MZ-like clusters was CXCR3 expression. CXCR3 is a crucial chemokine receptor involved in the homing of activated immune cells to sites of inflammation and in murine models of rheumatoid arthritis. CXCR3^+^ MZ-like B cells have been associated with T-bet expression and IL-10 production^45^. Nonetheless, the role of CXCR3 in MZ-like B-cell trafficking remains uncharacterized in humans, and further studies are needed to elucidate its contribution in pediatric LC. Lastly, pediatric LC was associated with increased expression of CD127, CD25 and PD1 across all B-cell populations except plasmablasts, which had lower CD25 and PD1 expression, suggesting reduced activation and, consequently, diminished antibody production in pediatric LC. Additionally, CD1c expression was decreased in B cells from CYP with LC. Reduced CD1c expression in B cells activated through the CD40-CD40L axis has been linked to impaired antigen presentation^46^, suggesting potential deficiencies in B cell–mediated immune responses in this population.

Based on our analysis of the B-cell compartment, we found that although the proportion of antibody responders and non-responders did not differ between the cohorts, CYP with LC had significantly lower levels of anti-RBD IgG and IgA antibodies. Conversely, there were no discernible differences in anti-S2 or anti-N IgG and IgA levels, indicating that CYP with LC may have a selectively diminished humoral response targeting the RBD region of the SARS-CoV-2 spike protein. Notably, these patterns contrast with findings in adult populations. Several studies have reported elevated antibody responses to the RBD and other Spike protein regions in adults with LC^19,31,47^. For example, Klein et al. reported increased levels of anti-spike, anti-S1 subunit, and anti-RBD IgG antibodies in adults with LC^19^. However, direct comparison with our study is limited, as the adult cohort had received two doses of a SARS-CoV-2 vaccine, whereas most children in our cohort were unvaccinated at the time of sample collection. Given the reduced anti-RBD antibody responses in CYP with LC, we also assessed neutralizing antibody activity.

Consistent with the decreased levels of anti-RBD IgG and IgA, neutralization assays revealed significantly lower SARS-CoV-2 neutralizing capacity in the LC cohort than in the controls. This is expected, as the RBD is a major target of neutralizing antibodies^48^. Interestingly, a recent study from the United Kingdom also reported reduced neutralization capacity in individuals with LC^49^. The reduced levels of anti-RBD antibodies and diminished neutralization capacity observed in pediatric LC may limit viral clearance during acute infection, enabling viral persistence in tissue reservoirs. This hypothesis is supported by several studies that identify low antibody titers during acute infection as a risk factor for LC^50–52^.

A notable limitation of our study concerns the size and composition of the control group, which comprised approximately one-fifth of the LC cohort. This imbalance resulted from recruitment constraints, as only children and adolescents without LC, inflammatory diseases or acute infections at the time of sample collection were eligible for inclusion. Consequently, we were limited to enrolling healthy pediatric volunteers whose parents/guardians provided consent for blood sampling. It is also important to acknowledge that at recruitment, COVID-19 vaccination had already begun in the pediatric population. Given these recruitment challenges, vaccination status could not be used as an exclusion criterion. As a result, most participants in the control cohort (72.2%) had been vaccinated prior to sample collection, compared to only 16.2% in the LC cohort. This discrepancy raises the possibility that the lower anti-RBD IgG and IgA levels observed in the LC cohort could be confounded by vaccination status. However, if vaccination were the primary driver of these differences, one would also expect reduced levels of anti-S2 antibodies, which were not observed. Moreover, a recent study reported no significant differences in total IgG levels between CYP who were vaccinated and those who experienced natural infection^53^. This observation supports the interpretation that the reduced anti-RBD IgG and IgA levels in CYP with LC are not solely attributable to differences in vaccination status but may instead reflect a distinct immunological profile associated with this condition. In terms of PBMCs immunophenotyping, previous research comparing individuals with prior SARS-CoV-2 infection to those who were vaccinated found no significant differences in the major T- and B-cell populations specific to SARS-CoV-2, suggesting that vaccination alone does not substantially alter the abundance of these immune subsets^54^. Importantly, all analyses in our study were adjusted for SARS-CoV-2 antigen exposure to control for its potential effect on both antibody titers and immune cell composition.

Another limitation concerns the timing of sample collection. Samples from the control cohort were obtained significantly closer to their most recent antigen exposure than those from the LC cohort. However, despite this difference, CYP with LC consistently demonstrated significantly lower levels of anti-RBD IgG and IgA antibodies, regardless of the time elapsed since their last antigen exposure. Moreover, SARS-CoV-2 neutralization capacity has been shown to remain stable for over six months after antigen exposure^55^, indicating that the differences observed in neutralization potential are unlikely to be due to sampling time.

Finally, using RF classification, we found that decreased CCR6 expression in monocytes was the most important feature that distinguished LC from controls. Remarkably, by incorporating only nine immunophenotyping variables, our model achieved almost 80% accuracy in identifying LC cases. Moreover, individuals with LC who were misclassified as controls had a higher clinical recovery rate than those who were correctly classified, suggesting a potential association between normalization of the immune profile and resolution of symptoms.

To conclude, our study provides a comprehensive characterization of the innate and adaptive immune responses in CYP with and without LC. The findings suggest that an inefficient innate immune response, characterized by a reduced capacity of APC to migrate to secondary lymphoid tissues and inflamed tissues, may result in diminished CD4 T cell responses. This, in turn, could lead to suboptimal germinal center reactions and, consequently, lower antibody levels and reduced SARS-CoV-2 neutralization capacity. Moreover, the observed activation of T, B and NK cells may reflect a compensatory mechanism in response to these diminished CD4 T-cell responses, whereby the limited pool of responding CD4 T cells becomes overactivated in an effort to maintain immune homeostasis.

## Author Contributors

JI-P, JM-P and SM-L were responsible for the design and execution of the study. AG-A, CC-A, MMe, CR and JD were responsible for the clinical data collection, confection of the cohort and sample collection. MN, MMa, JB and JC were responsible for the design of the spectral cytometry panel used. JI-P and SM-L were responsible for the spectral cytometry analysis. NP-L and JC were responsible for the specific SARS-CoV-2 antibody levels analysis. JI-P, TP, JB and BT were responsible for the specific SARS-CoV-2 neutralizing capacity analysis. VU, JI-P and SM-L were responsible for the statistical analysis. JI-P and SM-L were responsible for writing the manuscript. JI-P, SM-L and JM- P were responsible for the decision to submit the manuscript. NP-L, TP, MN, VU, FL, FM-L, JD, AG-A, CC-A, MMe, CR, MMa, JB, JC, BT and JM-P made important critical reviews and feedback on the manuscript. JM-P and SM-L as co-corresponding authors confirm that all authors have read and agreed with the current version of the manuscript. The co-corresponding authors had full access to all the data in the study and had final responsibility for the decision to submit for publication.

## Data Availability

Pseudonymized raw data and a data dictionary for each dataset are available from the corresponding authors upon request following the publication of this article. To gain access, data requesters will need to sign a data access agreement.

### Acknowledgments

Firstly, we thank all the participants and their families, without whom this study could not have been performed. We would also like to thank the IGTP Cytometry Core Facility and staff (MA Fernández Sanmartín and H Alzayat) for their contribution. We are grateful to the technical staff of IrsiCaixa for sample processing (L. Ruiz, E. Grau, R. Ayen, L. Gomez, C. Ramirez, M. Martinez, T Puig). We thank CERCA Programme/Generalitat de Catalunya for institutional support (2021 SGR 00452). Finally, we gratefully acknowledge all the data contributors, i.e., the authors and their originating laboratories for obtaining the specimens, and their submitting laboratories for generating the genetic sequence and metadata and sharing via the GISAID Initiative, on which the SARS-CoV- 2 variant period was retrieved.

## Statements and Declarations

The authors declare no competing interests. CR has received educational/consultancy fees from AstraZeneca, Sanofi and Merck Sharp & Dohme, all outside the submitted work. J.M-P. has received institutional grants and educational/consultancy fees from AbiVax, AstraZeneca, Gilead Sciences, Grifols, Janssen, Merck Sharp & Dohme and ViiV Healthcare, all outside the submitted work. S.M-L. has received institutional grants from Gilead Sciences outside the submitted work.

## Supplementary Tables and Figures Legends

Supplementary Figure 1: Manual gating strategy

Supplementary Figure 2: UMAP representing all clusters analyzed across the different cell populations. (A) Uniform manifold approximation and projection for dimension reduction (UMAP) representing the 17 clusters analyzed in the myeloid cell population using Flow-SOM and grouped depending on specific marker expression. (B) Uniform manifold approximation and projection for dimension reduction (UMAP) representing the 33 clusters analyzed in the NK-cell population using Flow-SOM and grouped depending on specific marker expression. (C) Uniform manifold approximation and projection for dimension reduction (UMAP) representing the 37 clusters analyzed in the T-cell population using Flow-SOM and grouped depending on specific marker expression. (D) Uniform manifold approximation and projection for dimension reduction (UMAP) representing the 31 clusters analyzed in the B-cell population using Flow-SOM and grouped depending on specific marker expression.

Supplementary Figure 3: Effect of time on antibody levels among CYP with and without LC. (A) Time to sample collection from the last exposure (weeks) in the LC and control cohort (median weeks [IQR]; LC: 20.3 [13.9-34.7]; controls: 11.6 [8.1-17.3]) (p<0.001). (B) Correlations of anti-RBD IgG (left) and anti-RBD IgA (right) in the LC and control cohorts (all p=n.s.) restricted to one-exposure samples. (C) Specific anti-RBD IgG (left) (p < 0.001) and IgA (right) (p = 0.028) antibody levels of responders. (D) Specific anti-S2 IgG (left) (p=n.s.) and IgA (right) (p=n.s.) antibody levels of responders. (E) Specific SARS-CoV-2 neutralizing capacity of responders in half-maximal inhibitory concentration (IC50) among the LC and control cohort (p = 0.004). Analyses of antibody and neutralization levels were restricted to one-exposure samples with no differences in time from the last exposure (median weeks [IQR]; LC: 13.9 [13.4-16.1]; controls: 11.0 [8.3-21.8]). Each dot represents an individual, and median and IQR values are indicated. p-values according to the Mann-Whitney test. Significant p-values < 0.05.

Supplementary Figure 4: Prevalence of responders and non-responders among CYP with and without LC. (A) Prevalence of anti-S2 IgG–specific responders and non-responders (%) among the LC and control cohorts. (B) Prevalence of anti-RBD IgG–specific responders and non-responders (%) among the LC and control cohorts. (C) Prevalence of Specific anti-N IgG responders and non-responders (%) among the LC and control cohorts. (D) Prevalence of Specific anti-S2 IgA responders and non-responders (%) among the LC and control cohorts. (E) Prevalence of specific anti-RBD IgA responders and non-responders (%) among the LC and control cohorts. (F) Prevalence of Specific anti-N IgA responders and non-responders (%) among the LC and control cohorts. (G) Prevalence of SARS-CoV-2–specific neutralizing capacity responders and non-responders (%) among the LC and control cohorts. LC n=99; controls n=18 in all analyses.

Supplementary Figure 5: Factor importance in random forest analysis. (A) Relative variable importance of the factors included in the random forest analysis.

**Supplementary Table 1:**
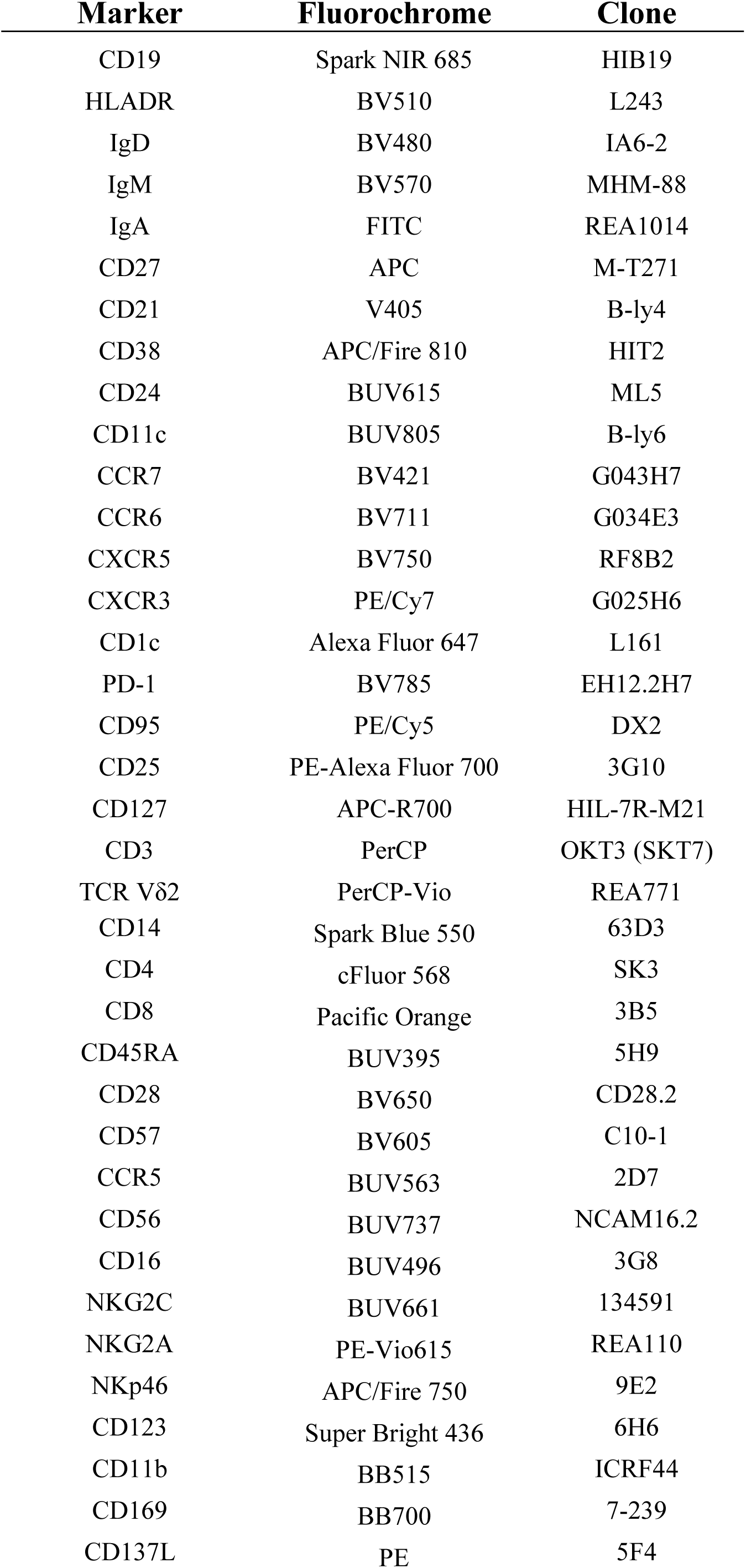
37-color panel used for spectral cytometry.

**Supplementary Table 2:** Cell count, marker annotation and myeloid cell population for 17 clusters

| Cluster | Cell count | Marker |  |  |  |  |  |  |  | Myeloid cell population 1 | Myeloid cell population 2 |
| --- | --- | --- | --- | --- | --- | --- | --- | --- | --- | --- | --- |
|  |  | CD14 | CD16 | CD11c | CD123 | CD1c | CD11b | HLADR | CD95 |  |  |
| 1 | 342,575 | + | + | + | - | - | + | + | + | Monocytes | Intermediate monocytes |
| 2 | 7,583,745 | + | - | + | - | - | + | + | + | Monocytes | Classical monocytes |
| 3 | 192,759 | - | + | + | - | - | - | + | + | Monocytes | Non-classical monocytes |
| 4 | 8,905 | - | + | - | - | - | low | low | + | Monocytes | Non-classical monocytes |
| 5 | 33,642 | - | + | - | - | - | low | - | + | Monocytes | Non-classical monocytes |
| 6 | 452,565 | - | - | + | - | + | - | + | + | Dendritic cells | cDCs |
| 7 | 68,514 | + | - | + | - | + | -/low | + | + | Dendritic cells | cDCs |
| 8 | 273,434 | + | - | + | - | - | low | + | + | Monocytes | Classical monocytes |
| 9 | 69,908 | - | - | - | - | - | - | low | + | Dendritic cells | CD123- CD11c- DCs |
| 10 | 74,789 | + | - | low | - | - | low | low | + | Monocytes | Classical monocytes |
| 11 | 11,384 | + | - | + | + | - | + | + | + | Monocytes | Classical monocytes |
| 12 | 1,441,128 | + | - | + | - | - | + | low | + | Monocytes | Classical monocytes |
| 13 | 649,848 | + | - | + | - | - | + | - | + | Monocytes | Classical monocytes |
| 14 | 653,258 | - | - | - | + | - | - | + | - | Dendritic cells | pDCs |
| 15 | 72,292 | - | - | + | - | - | - | + | - | Dendritic cells | cDCs |
| 16 | 354,045 | - | - | - | - | - | - | + | - | Dendritic cells | CD123- CD11c- DCs |
| 17 | 40,095 | + | - | - | - | - | + | + | -/+ | Monocytes | Classical monocytes |

**Supplementary Table 3:** Cell count, marker annotation and NK-cell population for 33 clusters

| Cluster | Cell count | Marker |  |  |  |  |  |  |  | NK-cell population |
| --- | --- | --- | --- | --- | --- | --- | --- | --- | --- | --- |
|  |  | CD56 | CD16 | CD57 | NKG2C | NKG2A | NKp46 | CCR5 | CXCR3 |  |
| 1 | 2,306,493 | dim | + | - | - | + | - | - | - | CD56dim |
| 2 | 46,356 | - | + | - | - | - | - | - | + | CD56- |
| 3 | 56,548 | dim | + | - | - | - | - | - | + | CD56dim |
| 4 | 393,957 | - | + | - | - | - | - | - | - | CD56- |
| 5 | 3,284,317 | dim | + | - | - | - | - | - | - | CD56dim |
| 6 | 18,331 | dim | + | - | - | - | - | + | + | CD56dim |
| 7 | 350,004 | dim | + | - | - | + | - | - | + | CD56dim |
| 8 | 33,772 | dim | - | - | - | + | - | - | + | CD56dim |
| 9 | 99,504 | dim | - | - | - | + | + | - | + | CD56dim |
| 10 | 187,417 | dim | + | - | - | + | + | - | + | CD56dim |
| 11 | 25,002 | - | + | - | - | - | + | - | low | CD56- |
| 12 | 150,315 | dim | + | - | - | - | + | - | + | CD56dim |
| 13 | 379,804 | dim | + | + | - | - | - | - | - | CD56dim |
| 14 | 85,325 | dim | + | - | - | + | + | + | + | CD56dim |
| 15 | 211,698 | bright | + | - | - | + | + | - | + | CD56bright |
| 16 | 45,591 | bright | - | - | - | - | + | - | + | CD56bright |
| 17 | 492,009 | bright | - | - | - | + | + | - | + | CD56bright |
| 18 | 42,988 | bright | + | - | + | - | + | - | + | CD56bright and NKG2C+ |
| 19 | 181,583 | dim | + | + | - | + | - | - | - | CD56dim |
| 20 | 273,266 | dim | + | + | + | - | - | - | - | CD56dim and NKG2C+ |
| 21 | 23,394 | dim | - | - | - | + | - | + | + | CD56dim |
| 22 | 17,574 | dim | - | - | - | + | + | + | + | CD56dim |
| 23 | 222,765 | dim | - | - | - | - | - | - | - | CD56dim |
| 24 | 17,905 | bright | - | - | + | - | + | - | + | CD56bright and NKG2C+ |
| 25 | 84,213 | bright | - | - | + | + | + | - | + | CD56bright and NKG2C+ |
| 26 | 46,549 | bright | - | - | - | + | - | - | low | CD56bright |
| 27 | 73,682 | bright | - | - | - | + | + | - | low | CD56bright |
| 28 | 327,552 | dim | + | - | + | - | - | - | - | CD56dim and NKG2C+ |
| 29 | 26,106 | dim | - | - | + | - | - | - | + | CD56dim and NKG2C+ |
| 30 | 6,280 | - | + | - | - | - | - | low | + | CD56- |
| 31 | 44,625 | dim | - | - | - | - | - | - | + | CD56dim |
| 32 | 42,584 | dim | - | - | - | - | - | + | low | CD56dim |
| 33 | 66,657 | dim | + | - | + | - | - | - | - | CD56dim and NKG2C+ |

**Supplementary Table 4:**
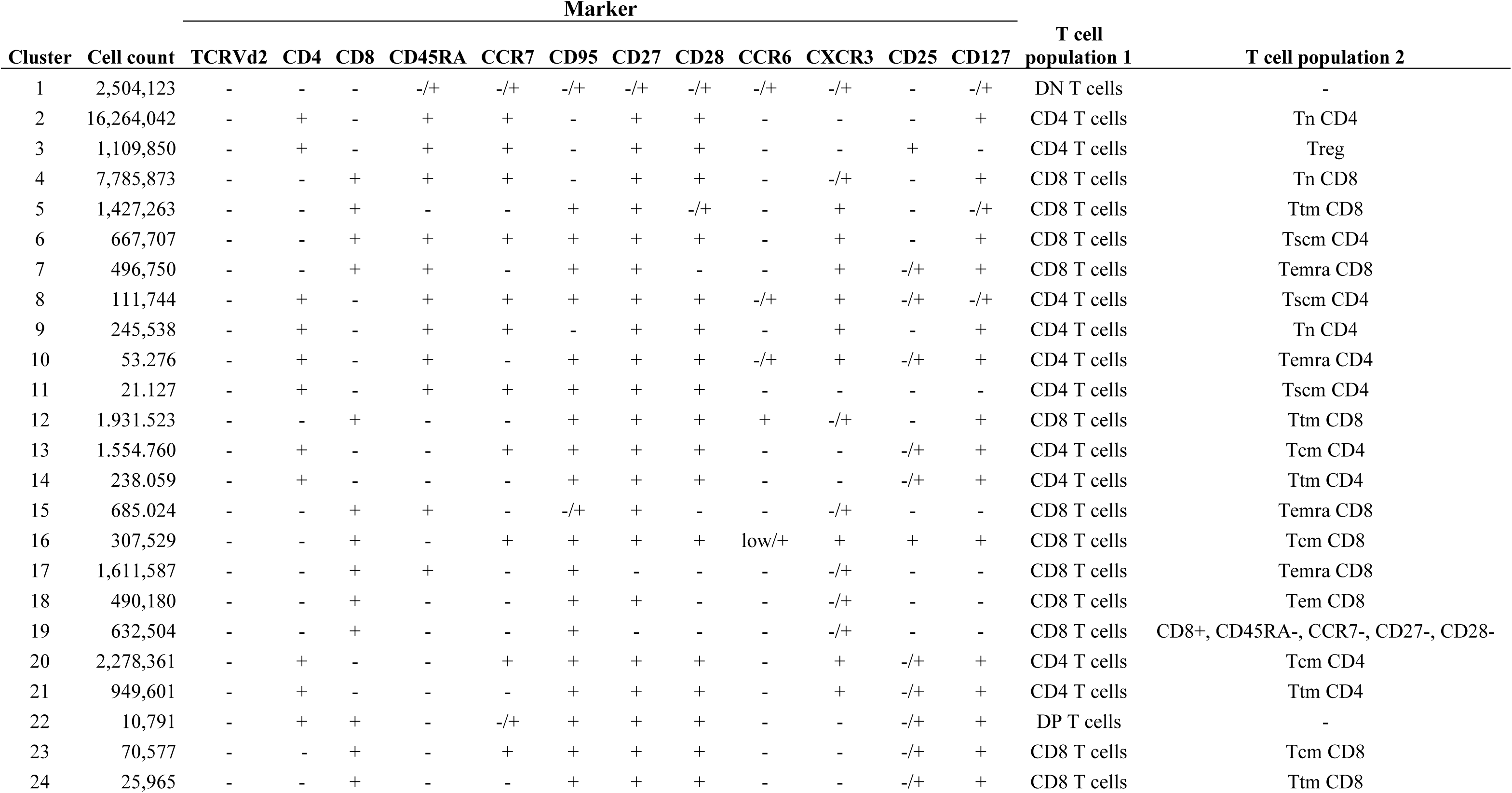
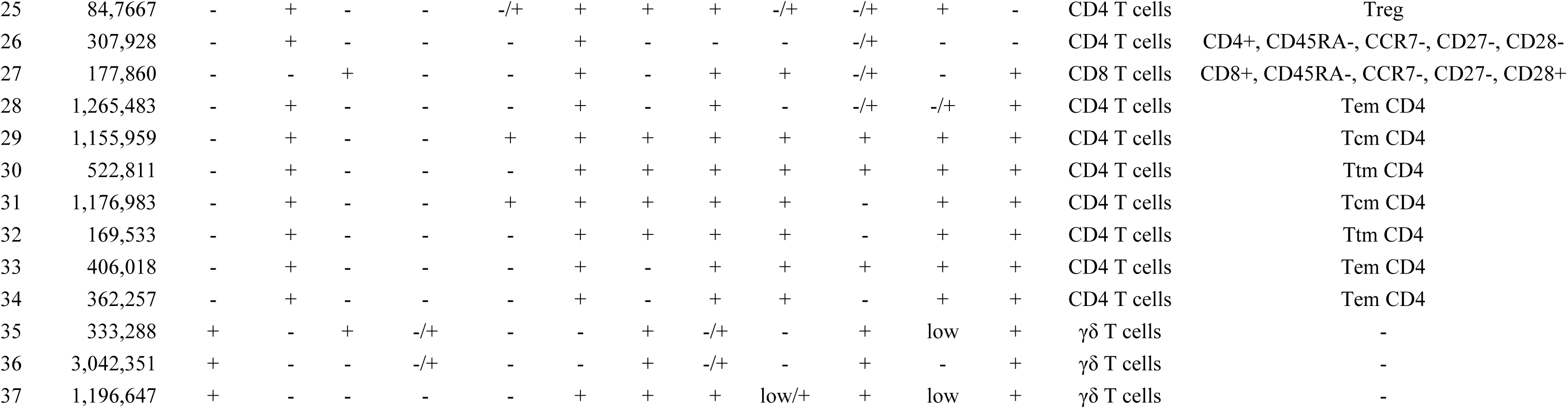
Cell count, marker annotation and T-cell population for 37 clusters

**Supplementary Table 5:** Cell count, marker annotation and B-cell population for 31 clusters

| Cluster | Cell count | Marker |  |  |  |  |  |  |  |  |  |  |  | B cell population |
| --- | --- | --- | --- | --- | --- | --- | --- | --- | --- | --- | --- | --- | --- | --- |
|  |  | IgD | IgM | IgA | CD27 | CD21 | CD38 | CD24 | CD11c | CCR7 | CCR6 | CXCR5 | CXCR3 |  |
| 1 | 113,849 | - | - | + | + | + | low | + | - | + | + | + | - | Switched memory B cells |
| 2 | 41,077 | - | - | + | + | + | low | low | -/+ | - | + | + | - | Switched memory B cells |
| 3 | 30,013 | - | -/+ | - | + | - | high | - | - | - | - | - | - | Plasmablasts |
| 4 | 27,104 | - | - | + | + | - | high | - | - | - | - | - | - | Plasmablasts |
| 5 | 29,758 | - | - | - | - | - | - | - | + | - | + | - | + | Switched memory B cells |
| 6 | 22,410 | - | - | -/+ | low | - | high | - | - | - | - | - | - | Plasmablasts |
| 7 | 3,089,971 | + | + | - | - | + | low | low | - | + | + | + | - | Naïve B cells |
| 8 | 19,772 | + | - | - | - | - | - | - | -/+ | - | + | - | + | IgD-only B cells |
| 9 | 50,271 | + | + | - | - | - | + | + | - | - | low | + | - | Transitional B cells |
| 10 | 175,072 | + | + | - | - | + | high | high | - | - | low | -/+ | - | Transitional B cells |
| 11 | 38,086 | - | - | + | + | + | low | + | - | + | + | + | + | Switched memory B cells |
| 12 | 69,857 | - | - | - | + | + | - | low | - | - | + | + | - | Switched memory B cells |
| 13 | 138,813 | - | - | - | + | + | low | + | - | + | + | + | + | Switched memory B cells |
| 14 | 52,186 | + | - | - | - | + | - | - | - | - | - | + | - | IgD-only B cells |
| 15 | 21,621 | - | - | - | - | - | - | - | - | - | - | low | - | Switched memory B cells |
| 16 | 28,476 | - | - | - | + | -/+ | low | low | -/+ | - | + | + | + | Switched memory B cells |
| 17 | 55,557 | - | - | - | + | + | - | high | - | - | + | + | - | Switched memory B cells |
| 18 | 206,392 | - | - | - | low | + | low | + | - | + | + | + | - | Switched memory B cells |
| 19 | 13,268 | - | -/+ | - | + | + | + | - | low | - | + | low | - | Switched memory B cells |
| 20 | 586,535 | + | + | - | - | + | + | + | - | + | + | + | - | Transitional B cells |
| 21 | 18,601 | - | + | - | + | + | - | high | - | - | + | + | + | IgM-only B cells |
| 22 | 20,245 | - | + | - | + | + | low | high | - | + | + | + | - | IgM-only B cells |
| 23 | 81,751 | + | + | - | + | + | - | high | - | - | + | + | - | MZ-like B cells |
| 24 | 338,391 | + | + | - | low | + | - | + | - | + | + | + | - | Naïve B cells |
| 25 | 25,560 | + | + | - | + | + | - | high | low | - | + | + | + | MZ-like B cells |
| 26 | 37,847 | + | + | - | low | + | - | + | - | + | + | + | + | Naïve B cells |
| 27 | 96,888 | + | + | - | - | + | low | low | - | + | + | + | + | Naïve B cells |
| 28 | 5,427 | + | - | - | + | + | - | -/+ | -/+ | - | + | + | -/+ | IgD-only B cells |
| 29 | 14,453 | + | + | - | - | low | - | - | -/+ | - | + | low | + | Naïve B cells |
| 30 | 20,324 | + | + | - | - | + | - | + | - | - | + | + | + | Naïve B cells |
| 31 | 14,056 | + | + | - | - | + | + | + | - | - | + | + | + | Transitional B cells |

**Supplementary Table 6:**
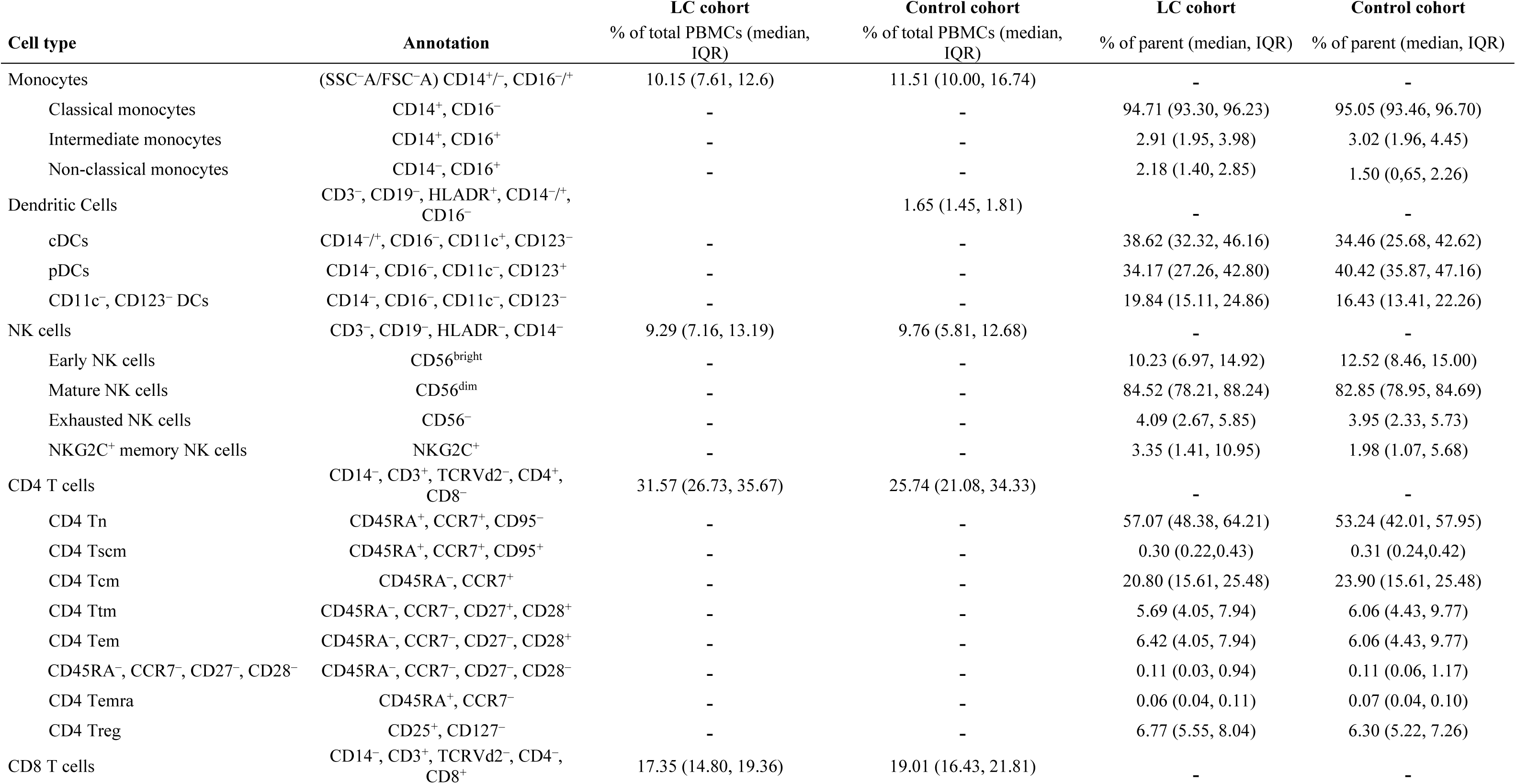
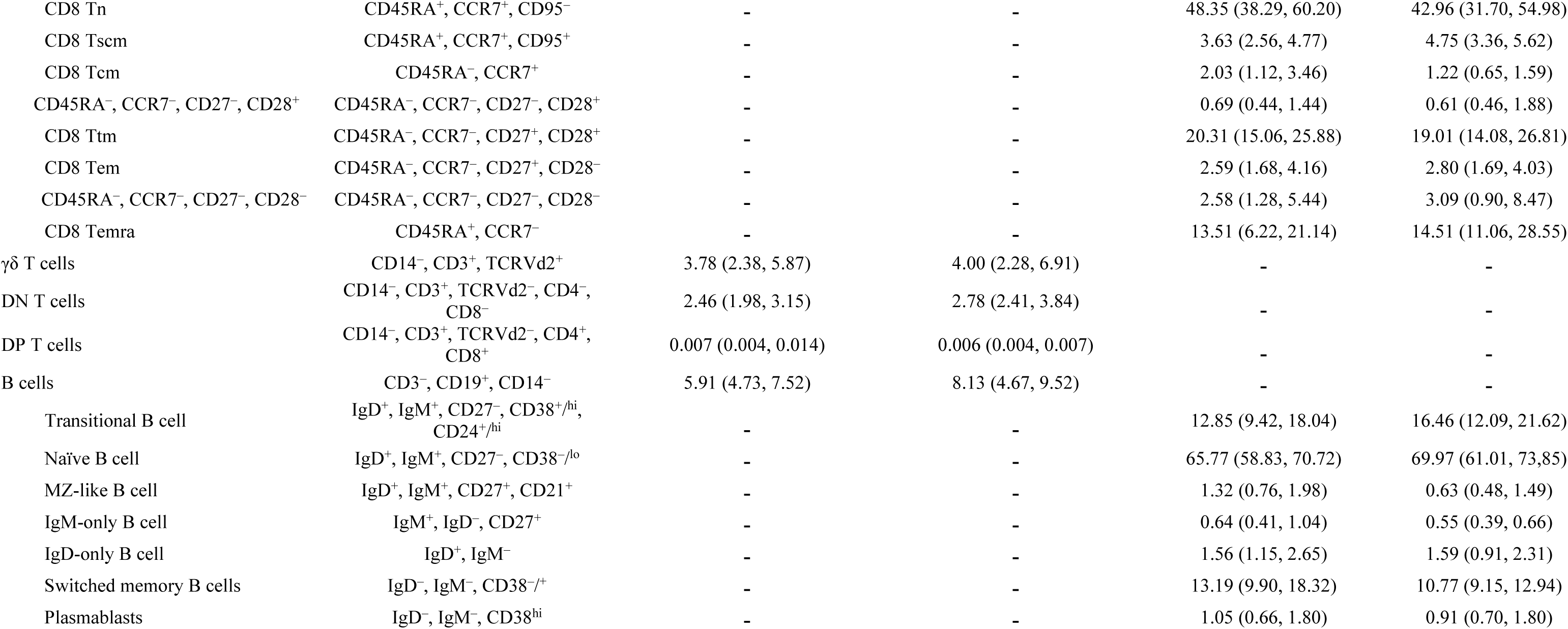
Frequency and annotation of different immune cell populations in CYP with and without LC.

**Supplementary Table 7:** Coefficients and p-values of linear regression models

|  | Coefficients |  |  |  | LRT p values |  |  |  |
| --- | --- | --- | --- | --- | --- | --- | --- | --- |
|  | Age | Sex=Male | Exposure= Two exposures | Group=PASC | Age | Sex | Exposure | Group |
| **S2 IgG** | 0.024 | -0.091 | 0.711 | 0.061 | 0.235 | 0.418 | 6.12E-07 | 0.669 |
| **RBD IgG** | 0.014 | 0.269 | 1.109 | -0.743 | 0.562 | 0.066 | 4.41E-10 | 8.99E-05 |
| **S2 IgA** | 0.039 | -0.001 | 0.318 | -0.061 | 0.038 | 0.925 | 0.011 | 0.652 |
| **RBD IgA** | 0.013 | 0.013 | 0.728 | -0.310 | 0.549 | 0.915 | 2.01E-06 | 0.046 |
| **Neutralization** | 0.039 | 0.204 | 0.568 | -0.330 | 0.192 | 0.112 | 1.47E-04 | 0.048 |

