## Supplementary figures and images for "Pediatric Long COVID Is Characterized by Myeloid CCR6 Suppression and Immune Dysregulation"

### Supplementary Figures 1-5

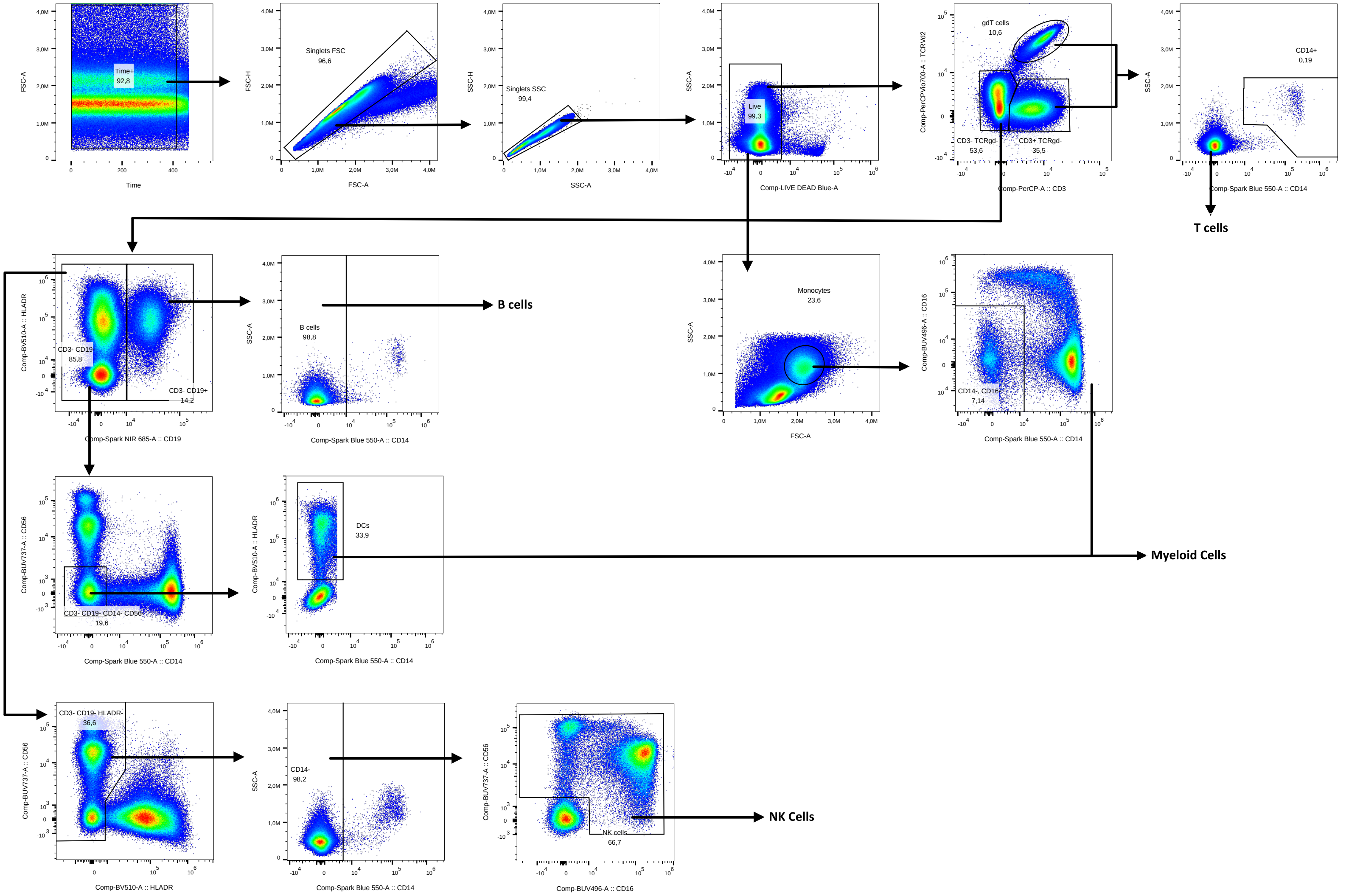

##### Supplementary Figure 2

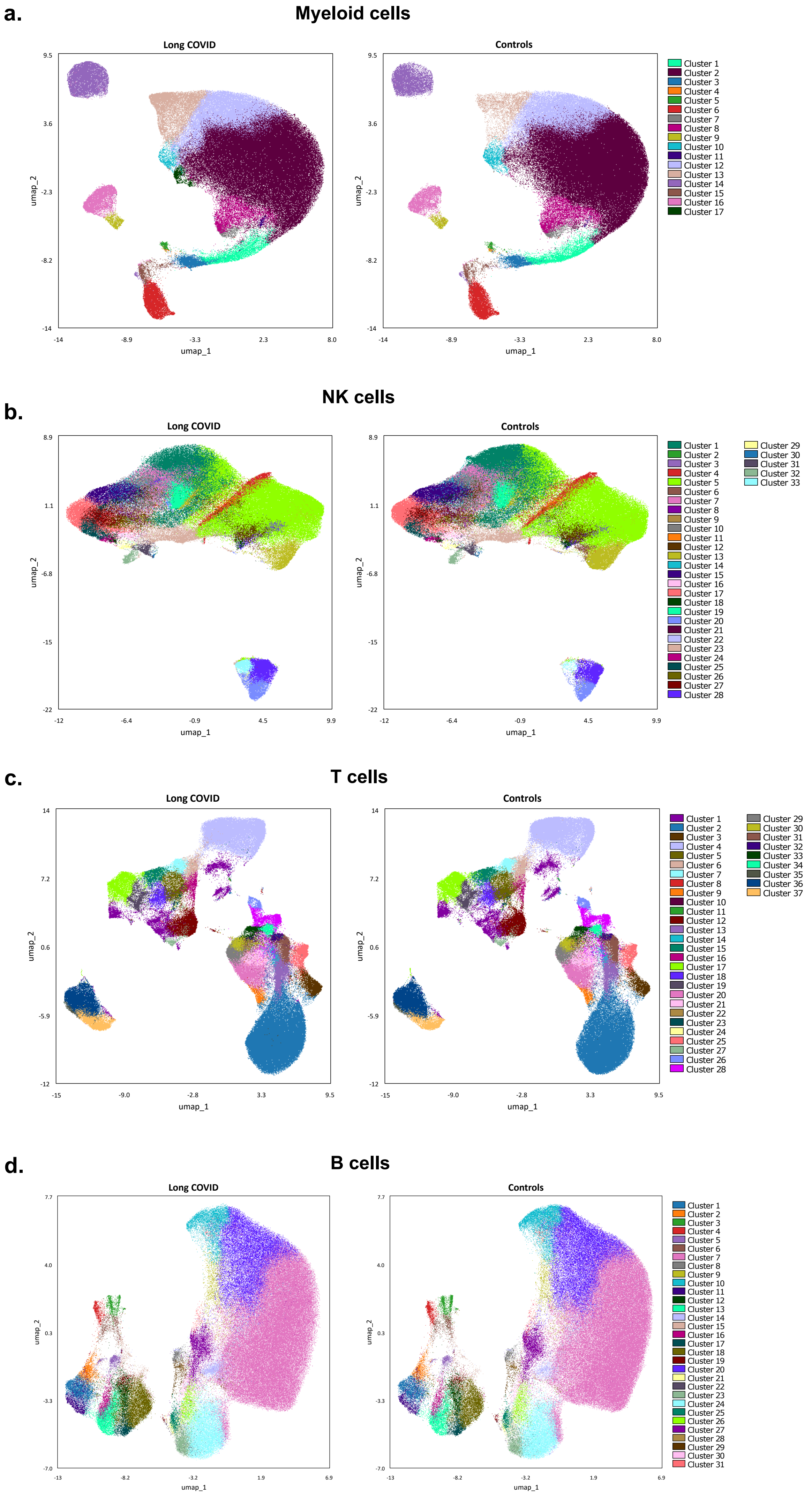

**Supplementary Figure 3**

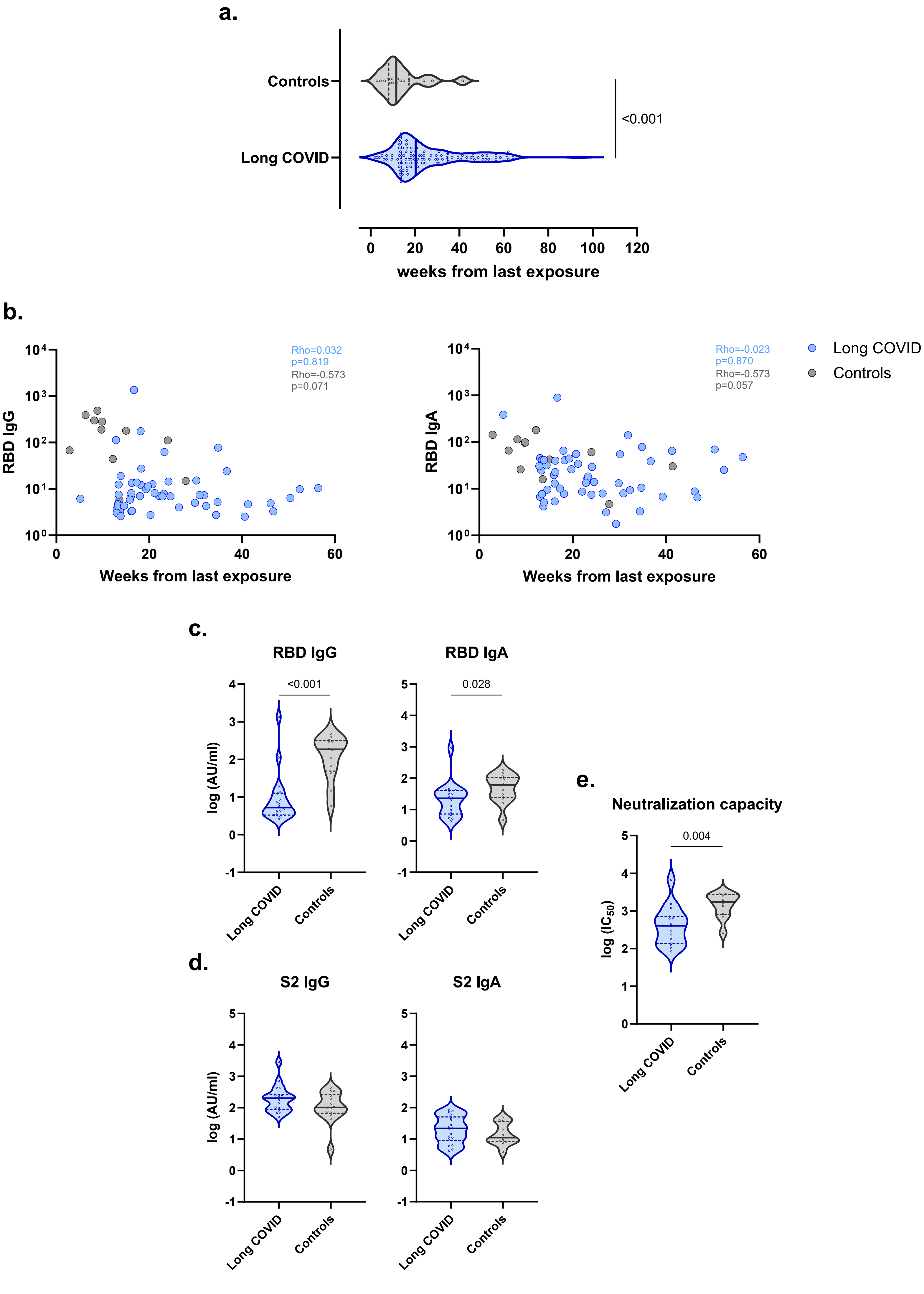

**Supplementary Figure 4**

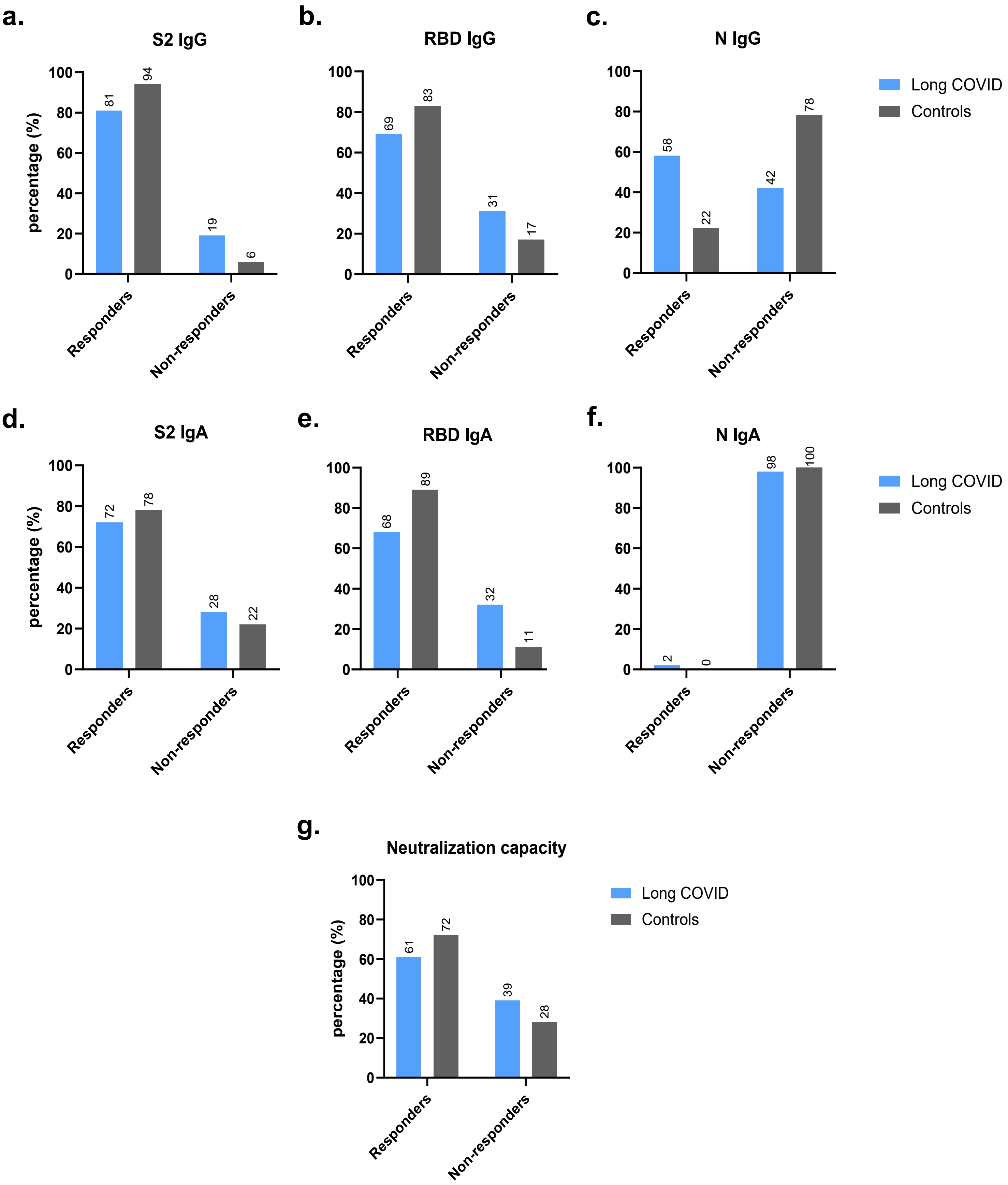

#### Supplementary Figure 5

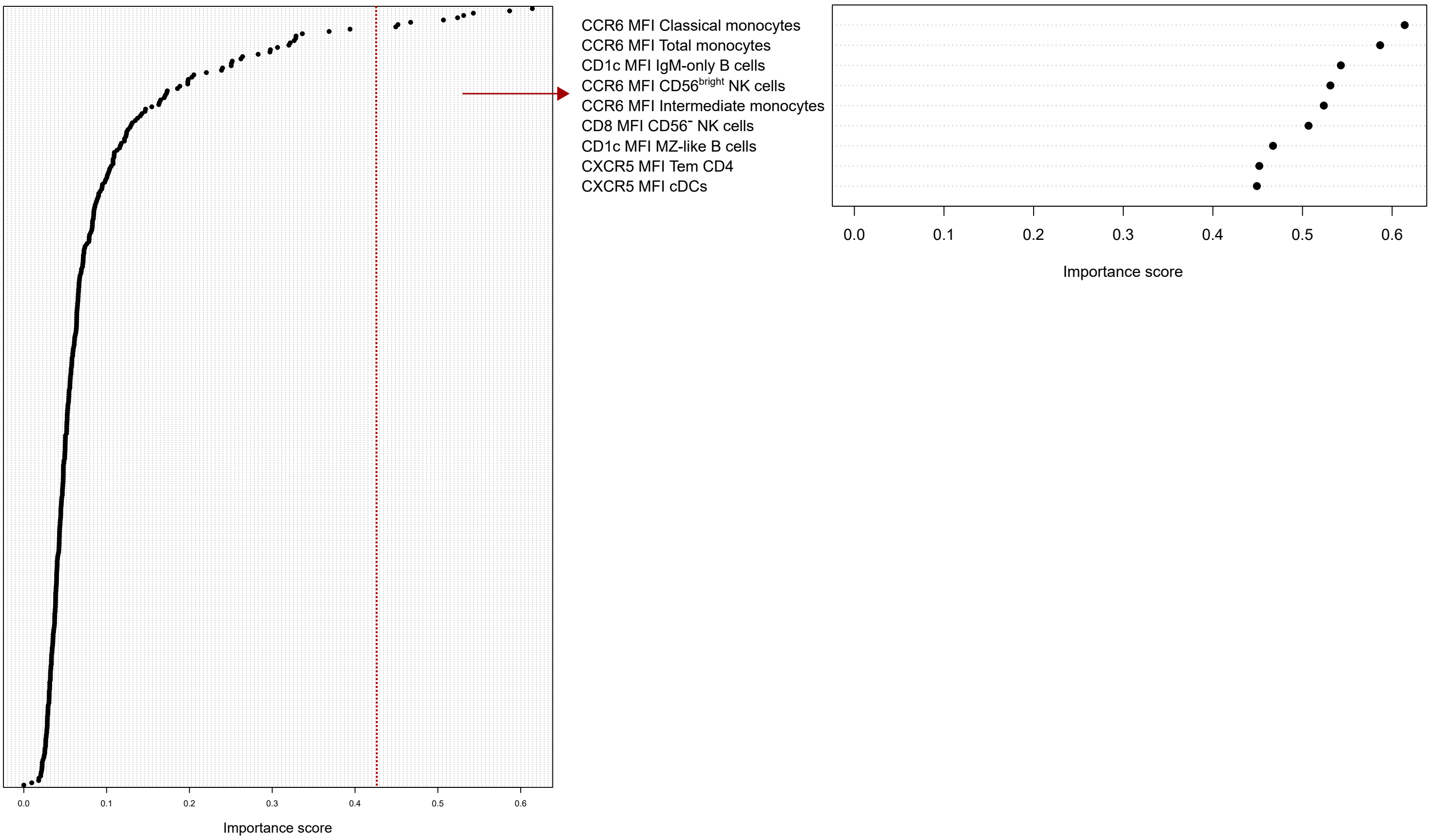
